# Subscaling of a cytosolic RNA binding protein governs cell size homeostasis in the multiple fission alga Chlamydomonas

**DOI:** 10.1101/2022.11.02.514835

**Authors:** Dianyi Liu, Cristina Lopez-Paz, Yubing Li, Xiaohong Zhuang, James G. Umen

## Abstract

Coordination of growth and division in eukaryotic cells is essential for populations of proliferating cells to maintain size homeostasis, but the underlying mechanisms that govern cell size have only been investigated in a few taxa. The green alga *Chlamydomonas reinhardtii* (Chlamydomonas) proliferates using a multiple fission cell cycle that involves a long G1 phase followed by a rapid series of successive S and M phases (S/M) that produces 2^n^ daughter cells. Two control points show cell-size dependence: Commitment in mid-G1 phase requires attainment of a minimum size to enable at least one mitotic division during S/M, and the S/M control point where mother cell size governs cell division number (n), ensuring that daughter distributions are uniform. *tny1* mutants pass Commitment at a smaller size than wild type and undergo extra divisions during S/M phase to produce small daughters, indicating that TNY1 functions to inhibit size-dependent cell cycle progression. *TNY1* encodes a cytosolic hnRNP A-related RNA binding protein and is produced once per cell cycle during S/M phase where it is apportioned to daughter cells, and then remains at constant absolute abundance as cells grow, a property known as subscaling (1). Altering the dosage of *TNY1* in heterozygous diploids or through overexpression increased Commitment cell size and daughter cell size, indicating that TNY1 is a limiting factor for both size control checkpoints. Epistasis placed *TNY1* function upstream of the retinoblastoma tumor suppressor complex (RBC) and one of its regulators, Cyclin-Dependent Kinase G1 (CDKG1) (2). Moreover, CDKG1 protein and mRNA were found to over-accumulate in *tny1* cells suggesting that CDKG1 may be a direct target of repression by TNY1. Our data expand the potential roles of subscaling proteins outside the nucleus and imply a control mechanism that ties TNY1 accumulation to pre-division mother cell size.

**Author Summary:** Size control is a fundamental property of cells which requires balancing cell growth with cell division, but the mechanisms used by cells to achieve this balance are only partly understood. The best-characterized mechanisms for size control to date involve fixed amounts of nuclear-DNA-bound inhibitory factors which repress cell division until cells grow past a minimum size threshold to overcome the inhibition. The unicellular green alga Chlamydomonas and many other algae and protists use a non-canonical cell cycle where cells can grow by many-fold in size before dividing, and then undergo multiple fission which involves successive rapid divisions to produce a uniform-sized population of daughters. In Chlamydomonas an unknown size homeostasis mechanism couples mother cell size to division number such that larger mother cells divide more times than smaller mother cells. Here, we identified and characterized a key factor governing size control in Chlamydomonas, a cytoplasmic RNA-binding protein and division inhibitor, TNY1, that is produced in a fixed amount in daughter cells and does not increase with cell growth, a property called sub-scaling. We found that TNY1 represses production of a cell cycle activator, CDKG1, during multiple fission to control daughter cell size. TNY1 is the first example of a cytosolic cell cycle inhibitor that does not depend on nuclear DNA binding to govern sub-scaling.

## Introduction

Size homeostasis is a fundamental property of proliferating cells and is achieved through mechanisms that balance cell growth rates with cell division rates. However, how cells sense and control size remains unexplored in most eukaryotic lineages. Two mechanisms for size homeostasis have been previously described: Adder-type mechanisms where a fixed mass is added in each cell cycle independently of birth size, and sizer mechanisms where one or more cell cycle transitions is dependent on cells reaching a minimum size (3, 4). Sizers have been characterized in several eukaryotes including budding yeast, mammalian tissue culture cells, and Arabidopsis meristems (3, 4). In each case, a titration mechanism operates where a cell cycle inhibitor is produced at a fixed absolute amount per cell in each cell cycle, a property known as subscaling, while an activator accumulates as cells grow (3, 5). At their critical size, cells have accumulated enough activator to overcome the inhibitor and allow cell cycle progression. The details of which proteins acts as the inhibitor or the activator differ in each species, but there are some systems-level similarities in several taxa including G1-S control with a nuclear-localized and/or chromatin associated factor as the subscaling inhibitor (1, 6).

Chromatin or nuclear DNA content is a naturally subscaled component of cells that has been exploited in Arabidopsis as a way of ensuring that the absolute amount of the inhibitor protein KRP4 apportioned to daughters is independent of birth size (7). In yeast and mammalian cells, chromatin-bound cell cycle inhibitor proteins, Whi5 and Rb respectively, are also subscaled and act as limiting inhibitors of cell cycle progression (8–10).

The unicellular green alga *Chlamydomonas reinhardtii* (Chlamydomonas) is a microbial model for plant cell cycles and for non-canonical multiple fission cell cycles that are used by many algae and other protists (11, 12). Multiple fission cell cycles partially uncouple cell growth and cell division: during a prolonged G1 phase, cells can grow more than ten-fold in size. Upon exiting G1, mother cells undergo (n) rapid alternating rounds of DNA synthesis and mitosis (S/M) and produce 2^n^ daughters within a common mother cell wall. Upon mitotic exit, the daughters hatch and enter either G0 or G1 phase due to nutrient availability (11, 13). The Chlamydomonas multiple fission cell cycle has two size control points or checkpoints. The Commitment point occurs in G1 phase, and is operationally defined by the transition from growth-dependence to growth-independence for completing at least one cycle of S/M. Cells must reach a minimum size to pass Commitment, and may continue to grow after Commitment for 5-7 hours, but this additional growth is optional for completing at least one cycle of S/M. Consequently, mother cells can begin S/M within a very large size range between two and twenty times the modal daughter size (11, 13). A second critical size checkpoint operates during the S/M phase and ensures that larger mother cells divide more times than smaller mother cells so that daughter sizes are in a uniform range regardless of the starting sizes of the mother cell population (11, 13). Thus, multiple fission incorporates a size control mechanism that is conceptually somewhat different than a simple gating mechanism used to control size in binary fission cell cycles.

Previous studies identified mutants that disrupted cell size homeostasis, including each subunit of the Chlamydomonas retinoblastoma tumor suppressor complex (RBC), MAT3/RBR, E2F1, and DP1 (14, 15). Interestingly, both Commitment size and the S/M size checkpoint were changed in these mutants (14, 15). Loss of function mutations in the *MAT3/RBR* gene caused cells to pass Commitment at a smaller size than wild type, and to divide too many times producing small daughters (14). In contrast, loss of function mutations in the *DP1* gene suppressed the *mat3/rbr* phenotype and caused cells to pass commitment at a larger size and to divide too few times leading to large daughters (15). Unlike the proposed model for size control in mammalian cells where the RB protein subscales (10), RBC subunits do not show this subscaling behavior in Chlamydomonas (16).

*cdkg1* was isolated in an insertional screen for size control defects. The mutant caused a large daughter cell phenotype and was found to act upstream of the RBC (2). CDKG1 encodes a D-cyclin dependent kinase (CDK) that phosphorylates the MAT3/RBR subunit of the RBC and is a limiting factor in mitotic size control. While loss of the protein in *cdkg1* mutants caused too few divisions and large cells, over-production of CDKG1 caused extra divisions leading to smaller daughter cells (2). CDKG1 protein is synthesized just before S/M begins with larger mother cells producing a higher nuclear concentration of CDKG1 than smaller mother cells. Nuclear CDKG1 concentration decreases with each round of cell division. Upon mitotic exit CDKG1 protein becomes undetectable and remains so until the S/M phase of the next cell cycle (2). It is unknown how *CDKG1* mRNA abundance and CDKG1 protein levels are modulated to control cell division number.

Here, we identified and characterized a Chlamydomonas heterogeneous nuclear ribonucleoprotein (hnRNP) related protein, TNY1, that acts as a cytosolic repressor in the size control pathway upstream of CDKG1 and the RBC. *tny1* mutants influenced Commitment and S/M size control and produced small daughters. TNY1 protein was produced once per cell cycle during S/M phase and apportioned to daughter cells where its absolute abundance stayed constant during G1 phase. Gene dosage alteration and over-expression experiments with *TNY1* both supported its role as a limiting regulator of mitotic size control. At least one key target of TNY1 repression is CDKG1, whose mRNA and protein abundance were negatively regulated by TNY1. TNY was found to be part of a ribonucleoprotein complex *in vivo*, and *in vitro* was able to bind the unusually long and uridine-rich 3’ untranslated region of the *CDKG1* mRNA. TNY1 is a novel example of a non-nuclear subscaling inhibitor which governs size control.

## Results

### TNY1 is a negative regulator of cell division upstream of CDKG1

*tny1-1* mutants were discovered in a forward insertional mutagenesis screen using a paromomycin antibiotic selection marker (paroR) with direct screening for size defects of plate-grown gametes using a Coulter Counter. *tny1-1* gametes showed a small size phenotype (Figure 1A) and the mutant was re-tested under more controlled vegetative growth conditions to assess daughter cell size. Wild-type parental strain CC-124 and *tny1-1* cultures were synchronized under a diurnal cycle and daughter cell sizes were measured. *tny1-1* daughter cells had a modal cell size of ∼50 µm^3^ compared with ∼80 µm^3^ for wild-type daughters (Figure 1B), with both strains passing Commitment and entering S/M with similar timing (Figure S1A), though with *tny1-1* populations always smaller than the control population when undergoing these two transitions (Figure S1B). The interval between Commitment and entering S/M was the same in wild type and *tny1-1* mutants, so the small size defects of *tny1-1* strains are not attributable to a shortened cell cycle duration (Figure S1A, B). We next generated populations of wild type and *tny1-1* mother cells and compared cell division numbers using a Commitment assay (Methods). When synchronized *tny1-1* and wild-type strains were sampled at the same time in late G1, division number profiles were similar (Figure S1C), despite the wild-type cells mother cells on average being 50-60% larger (Figure S1B); while in experiments where mother cell size distributions were matched between the two strains (modal size ∼230 µm^3^), *tny1-1* mother cells underwent an average of 2.8 rounds of multiple fission versus 1.4 rounds for wild type (Figure S1D). Together, these data show that while the overall timing of cell cycle events is normal in *tny1-1* mutants, the minimum Commitment size and mitotic control of *tny1-1* cells are both mis-regulated in a manner consistent with TNY1 acting as a negative regulator for size-dependent cell cycle control points.

**Figure 1.**
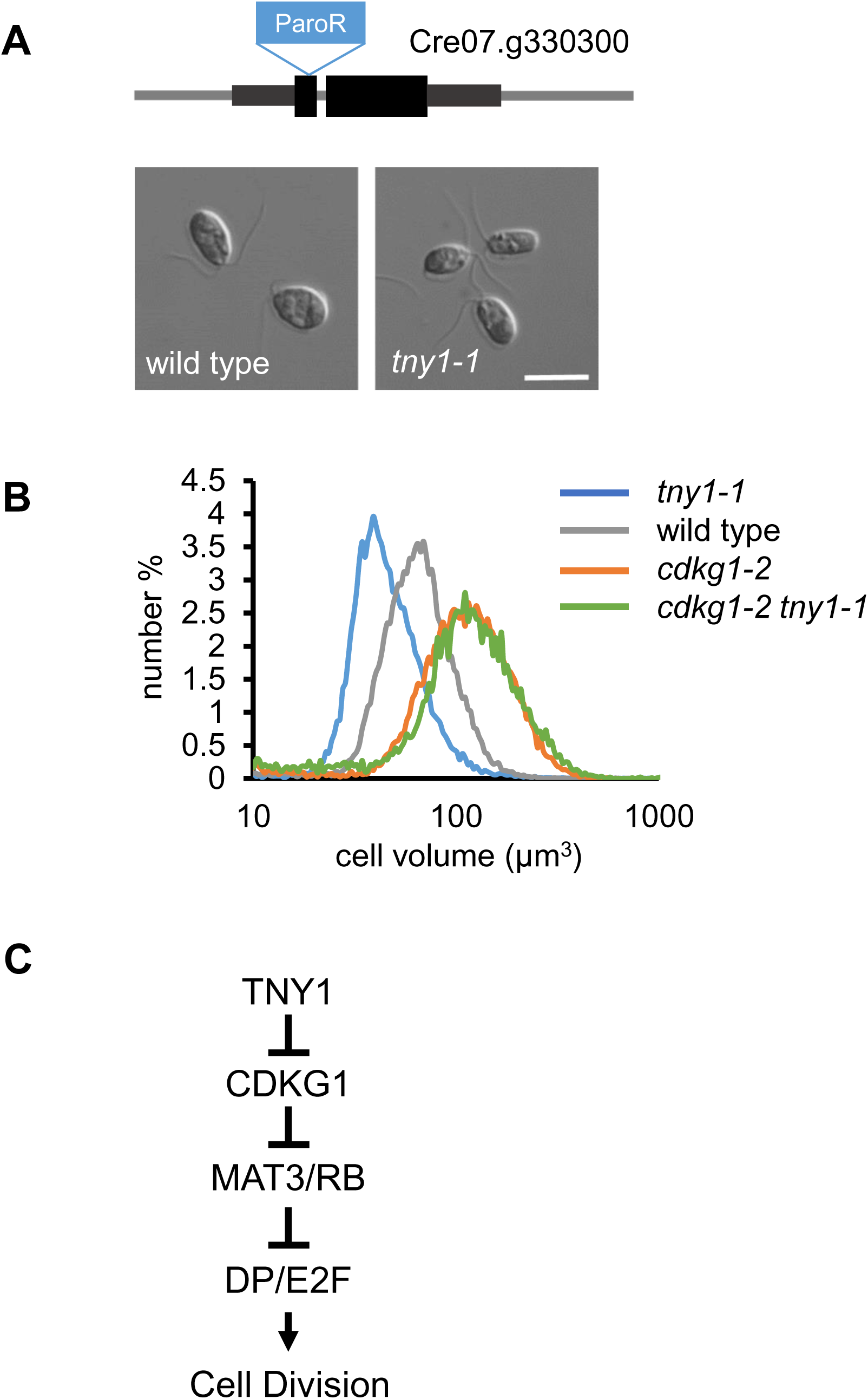
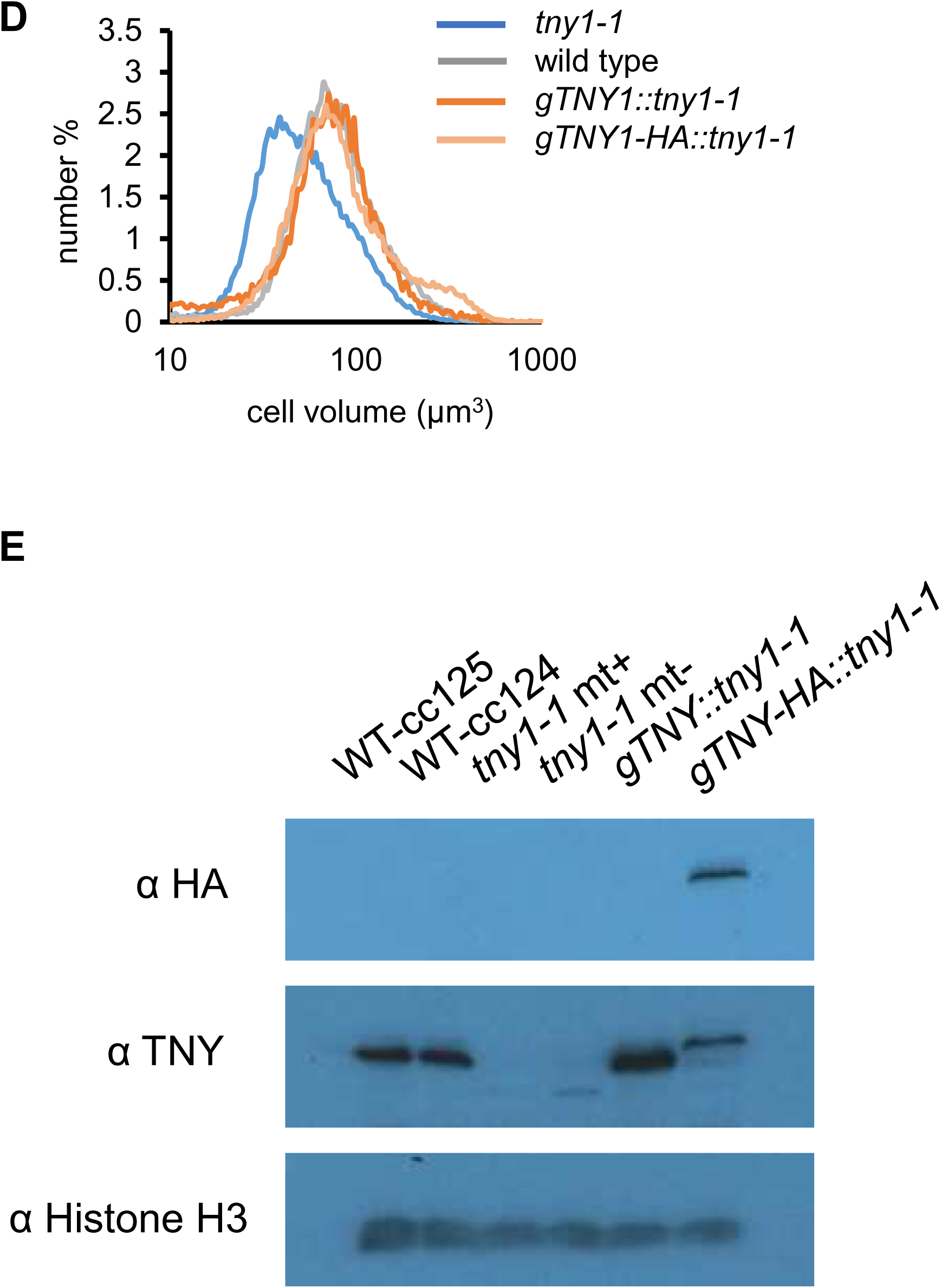
Identification of TNY1 as a regulator of cell size in the retinoblastoma pathway. (A) Upper panel, schematic of *TNY1* locus with location of an inserted paromomycin resistance marker (paroR in blue) in exon 1 that produced the *tny1-1* allele. Black rectangles, exons; dark gray rectangles, untranslated regions; narrow gray lines, introns, and intergenic regions. Lower panel, Differential Interference Contrast (DIC) images of daughter cells from wild-type parent strain CC124 and *tny1-1*. Scale bar = 10 µm. (B) Size distributions of daughter cells from *tny1-1*, wild type CC124, *cdkg1-2*, and *cdkg1-2 tny1-1*. Epistasis diagram showing positive (arrows) and negative (bars) regulators of size-dependent cell division. (C) *TNY1* functions upstream of *CDKG1.* (D) Size distributions of daughter cells from *tny1-1*, wild type CC124, and *tny1-1* rescued strains *gTNY1::tny1-1* and *gTNY1-HA::tny1-1*. (E) Immunoblot of SDS PAGE separated proteins from daughter cells of indicated genotypes using α-HA, α-TNY1, or α-histone H3 (loading control).

We next used epistasis experiments to determine the dependency *tny1-1* phenotypes on other size regulators. CDKG1 functions upstream of the RBC and *cdkg1-2* null mutants cause a large-cell phenotype. *cdkg1-2 tny1-1* double mutants had identical sizes as *cdkg1-2* single mutants indicating that TNY1 functions upstream of CDKG1 and the RBC, and does not appear to control cell size homeostasis through an independent mechanism (Figure 1B, C). Note that Commitment sizes for *cdkg1-2* and *cdkg1-2 tny1-1* (∼200 µm3) are very similar to the Commitment size (∼200 µm3) of a wild type strain (Figure S1E, F), indicating that *cdkg1-2* suppresses both the Commitment and the S/M size defects of *tny1-1*.

The *tny1-1* strain was found to contain a single insertion of the paroR marker in the first exon of Cre07.g330300 (17) (Figure 1A). *tny1-1* was back-crossed to wild type CC-125 and random progeny were chosen and scored for gamete cell size, mating type, and paromomycin resistance. The paroR segregants were small, while the paroS segregants were wild-type size indicating linkage between the paroR insertion and the *tny1-1* phenotype (Methods, Figure S1G). Rescue of the *tny1-1* small cell defect was performed by transforming constructs that contained either a full-length genomic fragment of wildtype Cre07.g330300 (gTNY1) or a version with a C-terminal triple hemagglutinin epitope tag (gTNY-3xHA). In both cases, normal daughter cell size was restored in transformants while no rescue was observed in control transformants bearing an empty vector (Figure 1D, Figure S1H, I). Rescue efficiency with either of two constructs was somewhat low (∼2%) but not atypical for Chlamydomonas rescues.

Immunoblotting of SDS-PAGE separated proteins from wild type, *tny1-1,* and rescued *tny1-1* strains using polyclonal antibodies raised against recombinant TNY1 protein or anti-HA antibodies detected proteins of the expected migration (∼48 kDa) in wild type and rescued strains showing that TNY1 expression was restored in those rescued lines (Figure 1E).

Together these experiments confirm that disruption of Cre07.g330300 causes the *tny1-1* phenotype.

### *TNY1* is predicted to encode a putative hnRNP A-related RNA binding protein

*TNY1* is predicted to encode a protein with two N-terminal RNA recognition motifs (RRMs) and a low complexity glycine-rich C-terminus (Figure 2A, Figure S2). This structure is found in eukaryotic heterogeneous nuclear ribonucleoproteins (hnRNPs) and other related RNA binding proteins that have diverse roles in nucleic acid regulation and metabolism, functioning as RNA or DNA binding proteins (18, 19). BLAST searching in different taxa was used to identify proteins related to TNY1 in animals, plants, and algae. These sequences were curated and used to estimate a maximum likelihood phylogeny which placed TNY1 in a clade of green algal TNY1-like homologs, and this TNY1 clade was sister to a larger grouping of plant tandem RRM hnRNP-like proteins suggesting a common origin at the base of the Viridaeplantae (Methods, Figure 2B). While Chlamydomonas encodes other hnRNP-like proteins, these grouped outside of the green algal TNY1 clade which appears to extend to near the base of the crown chlorophytes (Chlorophyceae/Trebuxiouphyciae/Ulvophyceae). No close matches to TNY1 were found in predicted proteomes of earlier diverging Chlorophyte branches in the Prasinophyte grade including *Micromonas* and *Ostreococcus* which both have reduced genomes and may have lost the ancestral *TNY1*-related genes.

**Figure 2.**
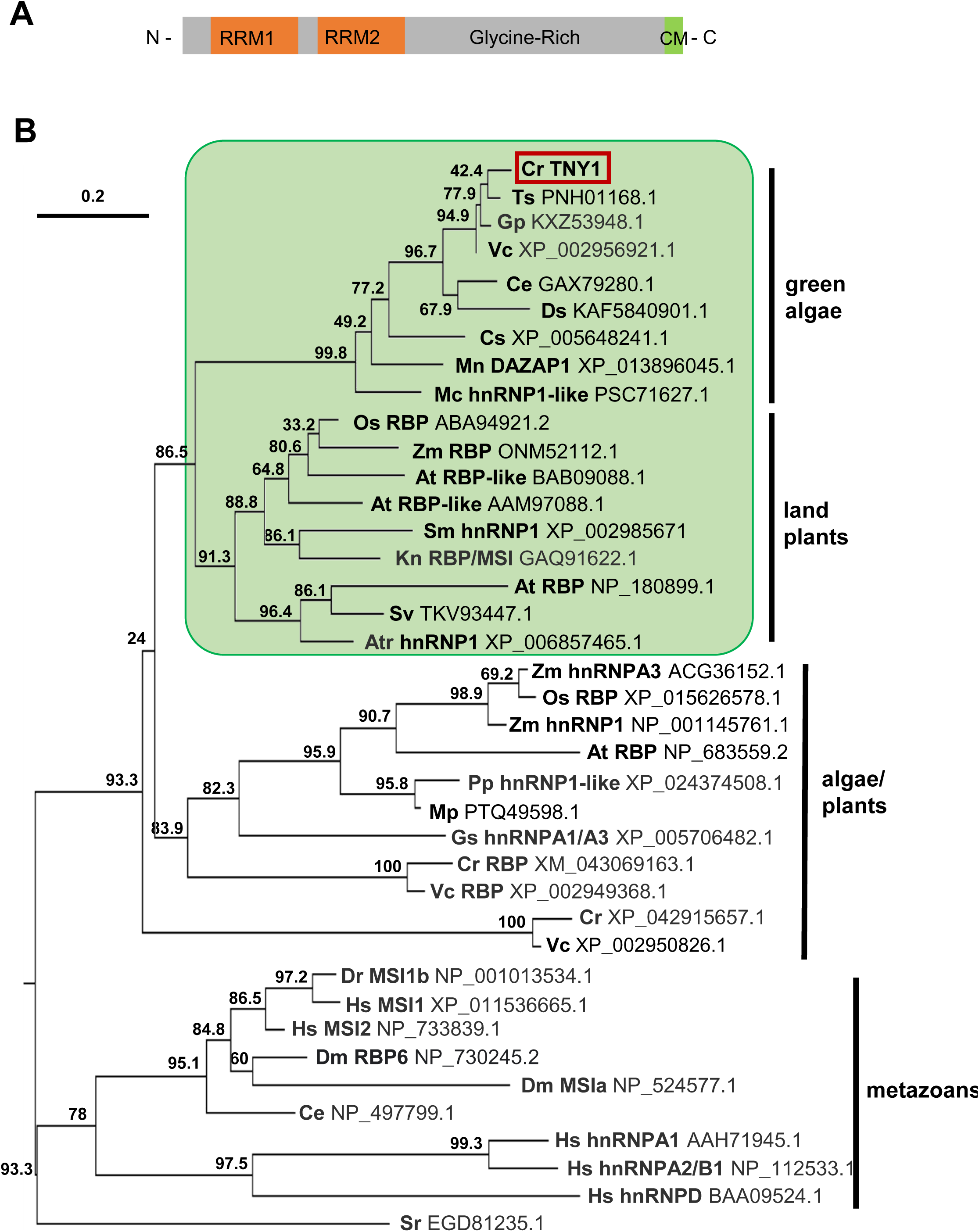
*TNY1* encodes a hnRNP-related RNA binding protein. (A) Schematic of predicted TNY1 protein domain structure from N to C terminus. Two RNA binding motifs (RRM1 and RRM2, orange bars) are followed by a glycine-rich region and a short conserved motif (CM) at the C-terminus. (B) Maximum likelihood phylogeny TNY1 and related hnRNP related proteins in indicated taxonomic groups. Species abbreviations are followed by protein names and NCBI protein IDs. Cr, *Chlamydomonas reinhardtii*. Ts, *Tetrabaena socialis*. Gp, *Gonium pectorale*. Vc, *Volvox carteri*. Ce, *Chlamydomonas eustigma*. Ds, *Dunaliella salina*. Cs, *Coccomyxa subellipsoidea*. Mn, *Monoraphidium neglectum*. Mc, *Micractinium conductrix*. Os, *Oryza sativa*. Zm, *Zea mays*. At, *Arabidopsis thaliana*. Sm, *Selaginella moellendorffii*. Kn, *Klebsormidium nitens*. Sv, *Setaria viridis*. Atr, *Amborella trichopoda*. Pp, *Physcomitrella patens*. Mp, *Marchantia polymorpha*. Gs, *Galdieria sulphuraria*. Dr, *Danio rerio*. Dm, *Drosophila melanogaster*. Ce, *Caenorhabditis elegans*. Hs, *Homo sapiens*. Sr, *Salpingoeca rosetta*.

### TNY1 is localized in the cytosol

To determine the subcellular localization of TNY1, a genomic *TNY1* construct with a C-terminal fusion of *mCherry* (20) was used to rescue *tny1-1* mutant cells and generate *gTNY1-mCherry::tny1-1* strains with fusion protein expression confirmed by immunoblotting (Figure S3). Live cell confocal fluorescence microscopy revealed that TNY1-mCherry is detectable in the cytosol throughout the vegetative cell cycle with a weak but significant signal detected at all stages (Figure 3A). Indirect immunofluorescence using anti-HA antibodies targeting tagged TNY1-HA confirmed the cytosolic location and showed exclusion of TNY1 protein signal from the nucleus (Figure 3B).

**Figure 3.**
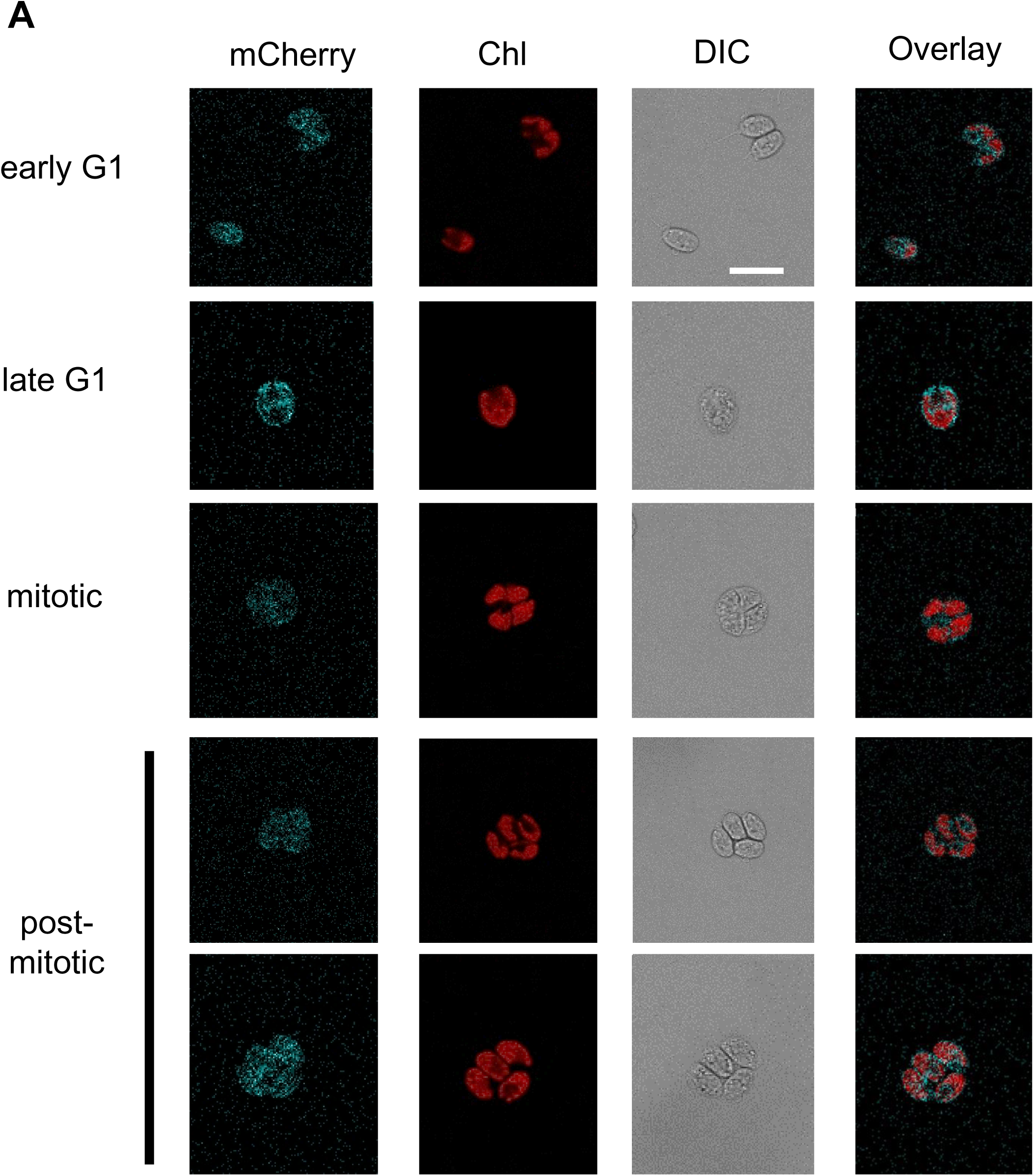
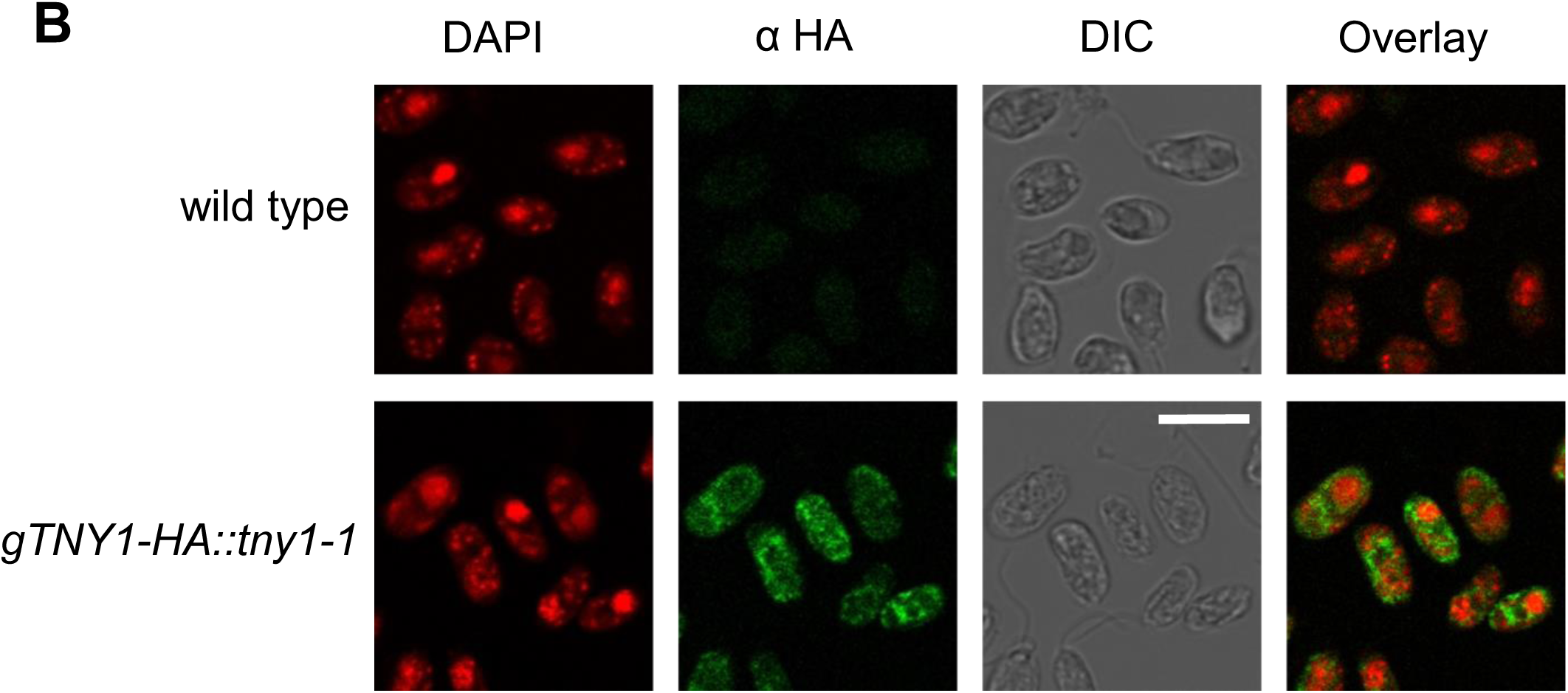
TNY1 is localized in the cytosol. (A) Brightfield and confocal fluorescence images of live cells at different cell cycle phases left side labels) expressing a functional TNY1-mCherry fusion protein. mCherry signal is false colored cyan and chlorophyll fluorescence (Chl) is false colored red. Merged fluorescent images are in the left column and brightfield images in the right column. Scale bar = 10 μm. (B) DIC and widefield immunofluorescence microscopy images of wild type (CC124) and *gTNY1-HA::tny1-1*. Daughter cells were fixed and immuno-stained for HA epitope (green). DAPI staining (red) was used to visualize nuclei. Merged fluorescence image is on the right. Scale bar = 10 μm.

### TNY1 regulation and subscaling throughout the cell cycle

To determine the accumulation pattern of *TNY1* mRNA during the cell cycle wild-type cultures were synchronized under a standard diurnal cycle (12hr:12hr light:dark) and RNA samples were prepared from cells at different time points and used for quantitative RT-PCR. *TNY1* mRNA was present at very low levels during G1 phase and rose sharply to a peak toward the middle/end of S/M phase, and then declined slowly in the dark phase after division (Figure 4A top panel). This experiment largely reproduced the results of (21) and (22), where the timing of *TNY1* mRNA accumulation coincided with that of many late mitotic and cilia-related genes.

**Figure 4.**
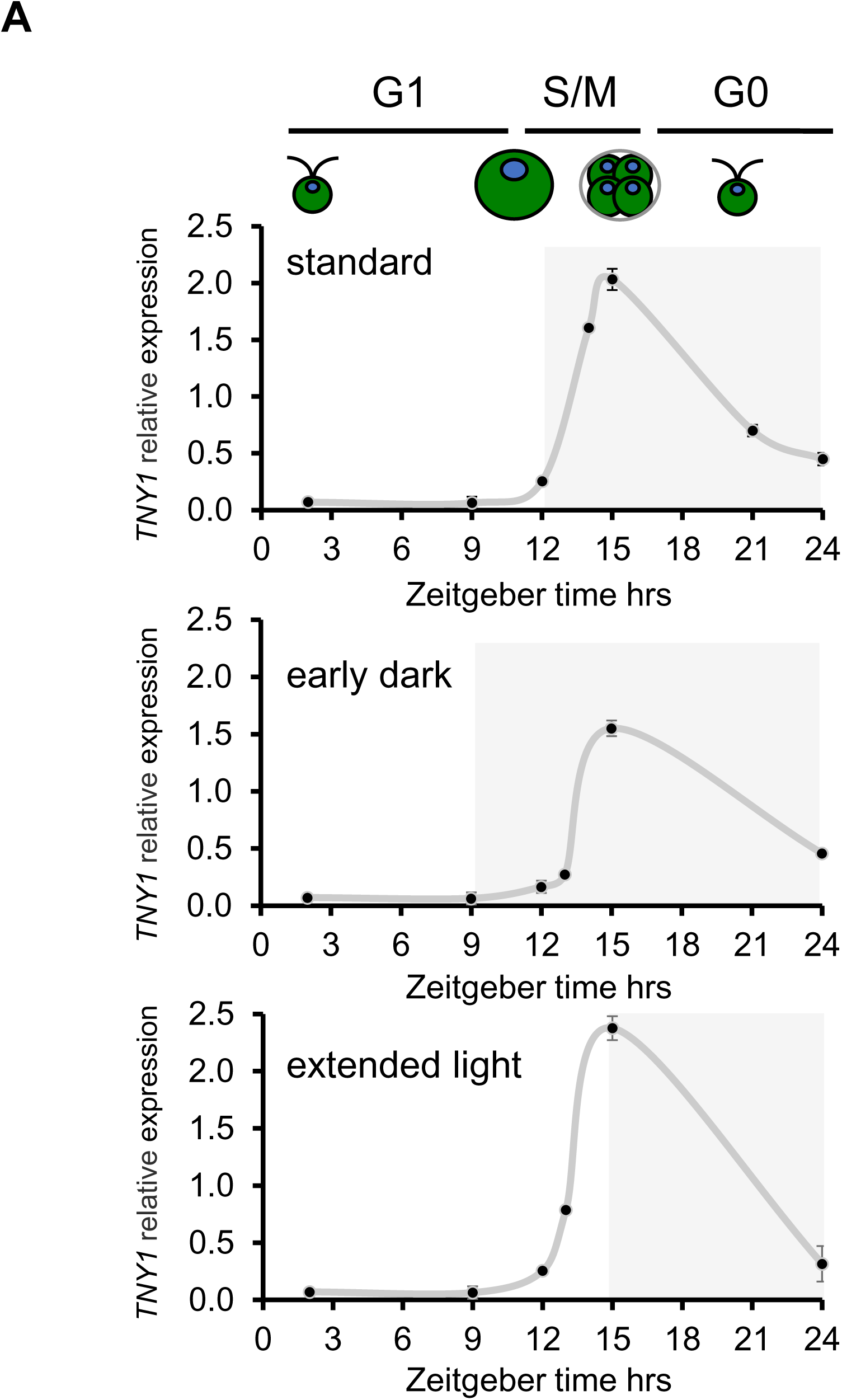
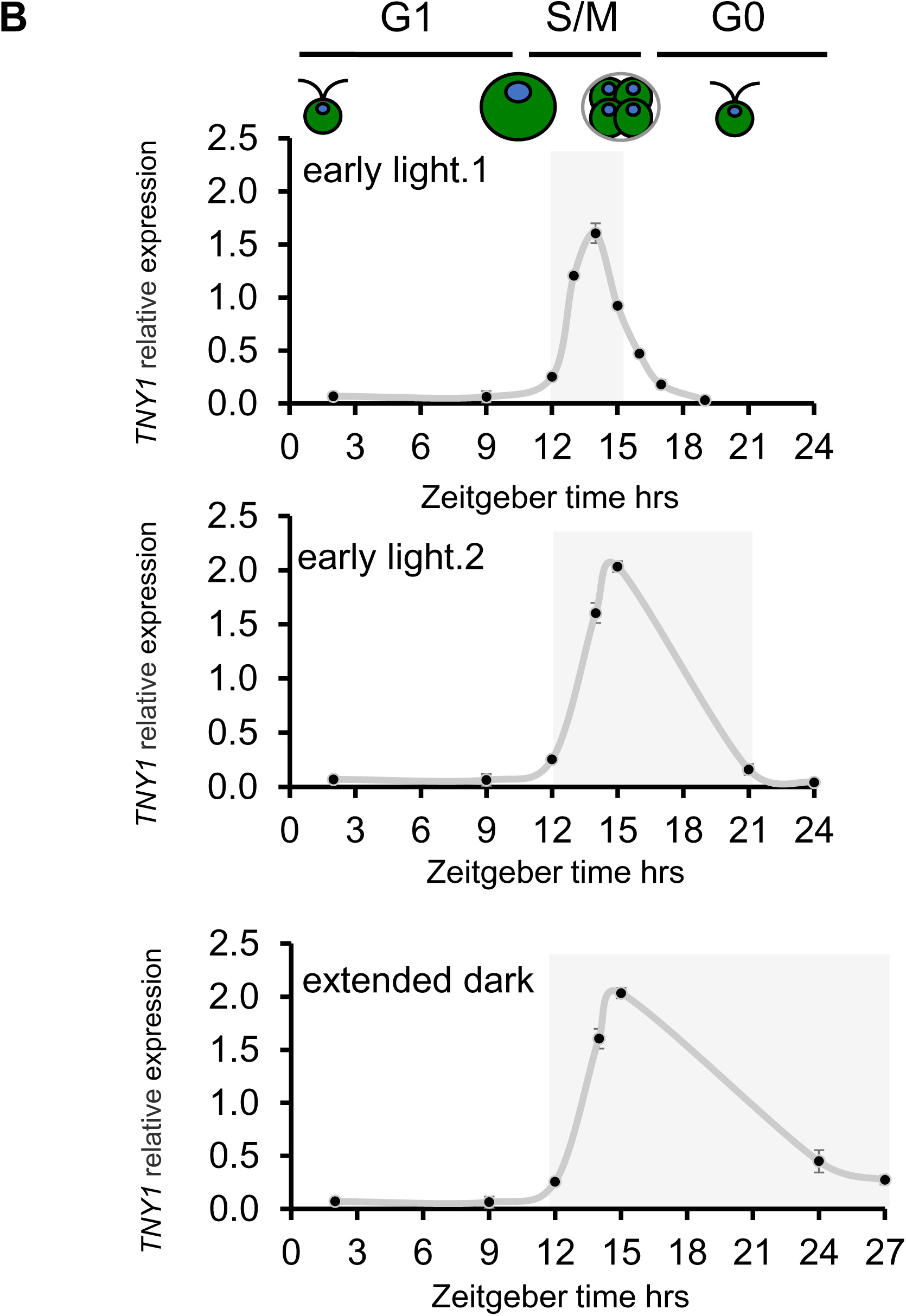
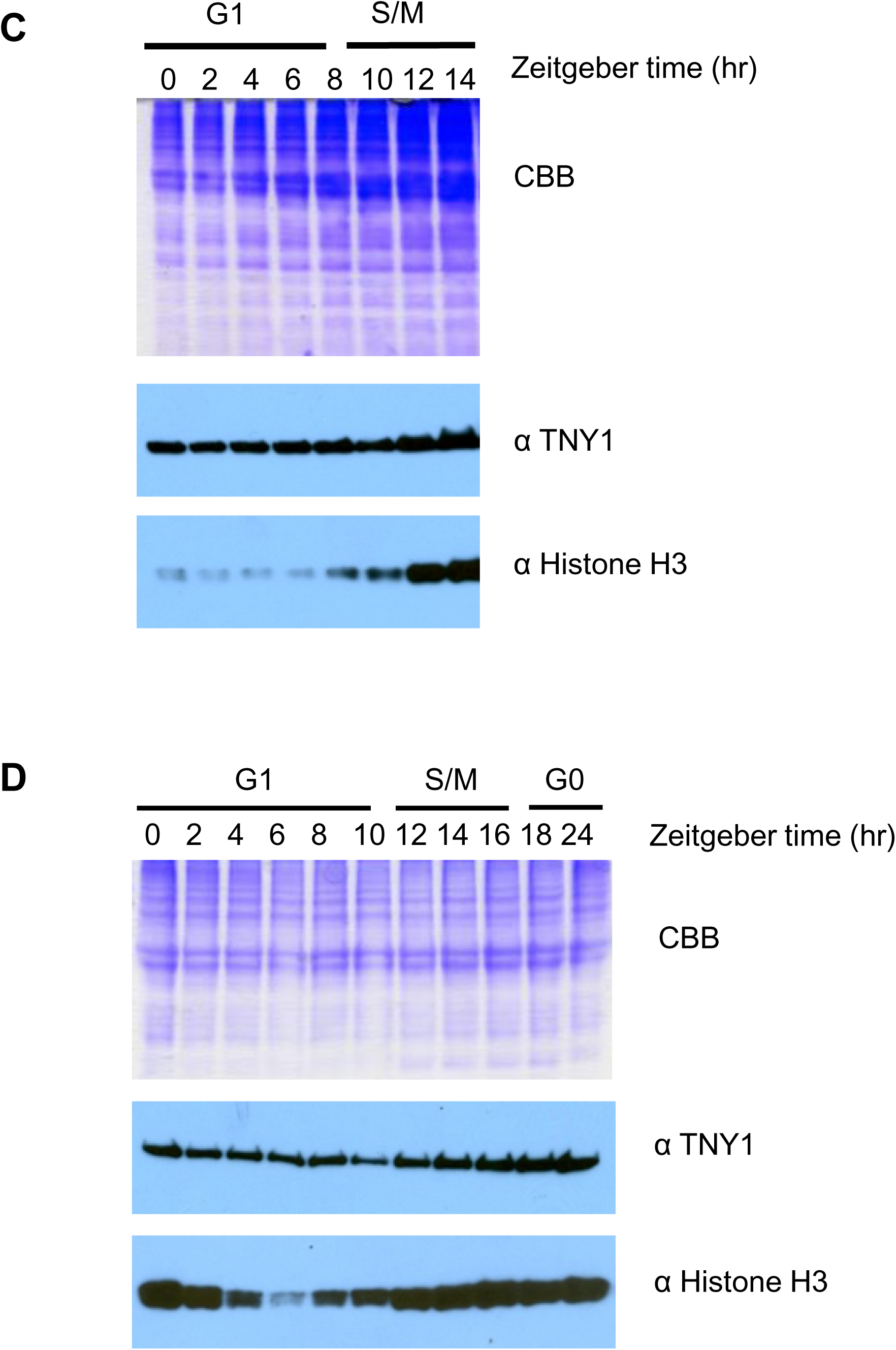
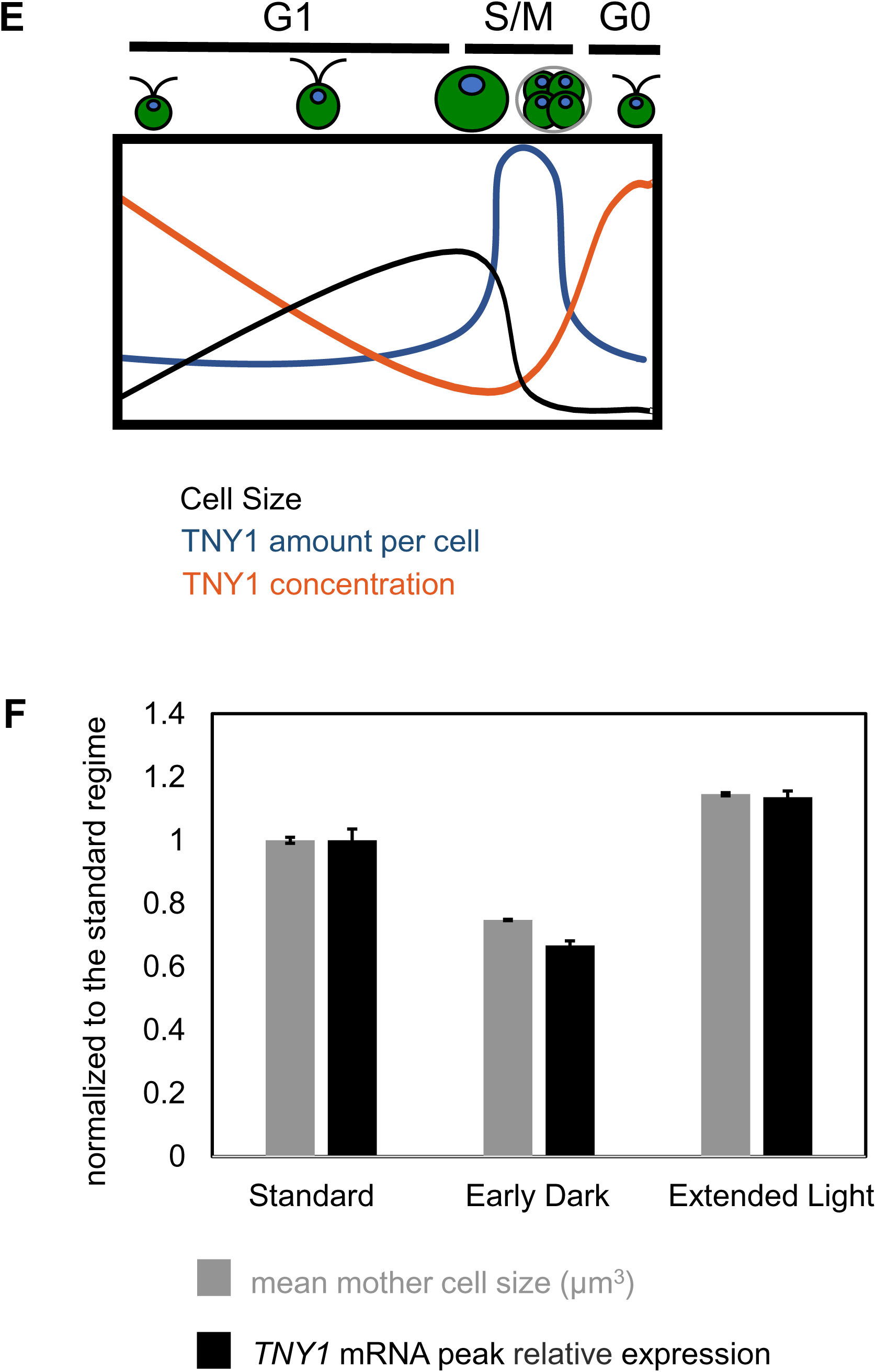
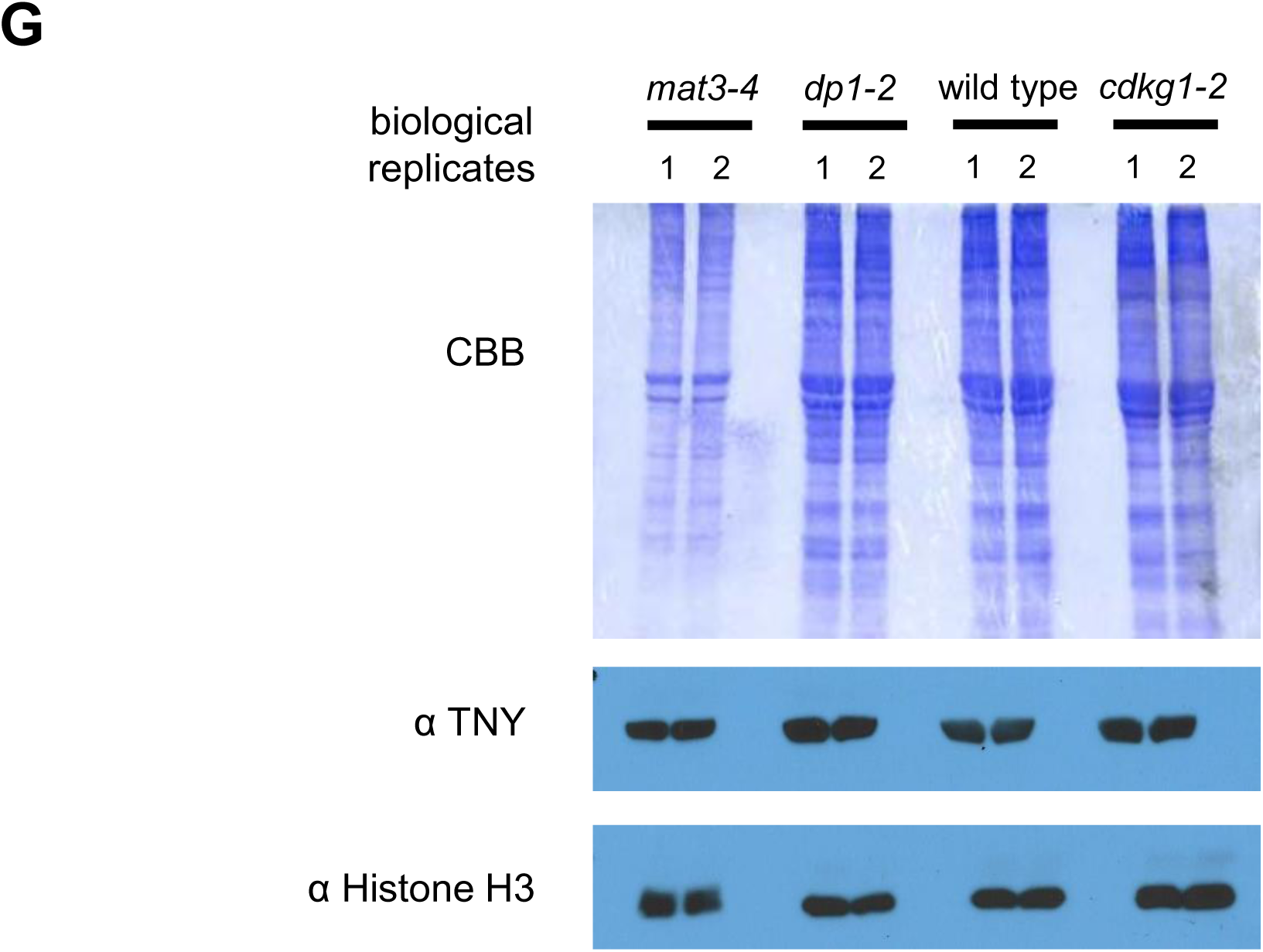
Cell cycle control of *TNY1* mRNA and TNY1 protein accumulation. (A) qRT-PCR data time series for *TNY1* mRNA accumulation in synchronous cultures with light and dark phases shown in white or gray, respectively, and cell cycle phasing cartooned above. The middle and bottom graphs show data for cultures synchronized under standard conditions (top) and released into a modified regime of early dark (middle panel) or extended light (bottom panel). All data were normalized against control transcript for *GBLP*. Error bars: high and low values of two biological replicates. The value of each biological replicate is calculated as the average of two technical replicates. Top panel, standard regime - 12hr:12hr light:dark. Middle panel, early dark - 15hr:9hr light:dark. Bottom panel, extended light - 9hr:15hr light:dark. Under all three different diurnal regimes, most of the cells start to divide at ZT 12 and finish division at ZT 15. (B) qRT-PCR data time series for *TNY1* mRNA accumulation in cultures similar to (A) but with altered lengths of the final dark period. Top panel, early light.1 with lights on at ZT 15. Middle panel, early light.2 with lights on at ZT 18. Bottom panel, extended dark through ZT 27. (C) and (D) Immunoblots of whole cell lysates from synchronized standard cultures shown in panel (A) Upper panel were fractionated on SDS PAGE gels and probed with either α-TNY1 or α-histone H3 as a control. Upper panel shows Coomassie staining and lower two panels show immunoblots with indicated antibodies. Gels were loaded with equal cell numbers per lane. (C) or equal protein per lane (D). (E) Schematic of TNY1 protein subscaling behavior showing how total protein per cell and concentration change during the cell cycle. The spike in TNY1 per cell during division can be interpreted as a sum of its accumulation in postmitotic mother cells with unhatched daughters. (F) TNY1 mRNA quantitation in mitotic cultures with same numbers of mother cells that are different mean sizes due to shortened or lengthened final light periods with data normalized to the standard 12:12 L:D regime. (G) Whole cell lysates of daughter cells from indicated genotypes (with two replicates 1 and 2) were loaded with equal cell number per lane, fractionated by SDS-PAGE and immunoblotted using α-TNY1 or α-Histone H3. Coomassie blue staining is shown.

The trigger for *TNY1* mRNA accumulation is likely to be cell division, but we could not rule out diurnal control or the light-to-dark transition as the drivers of *TNY1* expression. To distinguish these possibilities we used two alternative diurnal regimes where peak S/M phase (12-14 hrs ZT) did not coincide with the light-to-dark transition (early dark regime - 15hr:9hr light:dark; extended light regime - 9hr:15hr light:dark). In both alternative regimes, *TNY1* mRNA peaked with S/M phase and was not significantly shifted by the timing of the light-dark transition (Figure 4A).

*TNY1* mRNA declines gradually after S/M and becomes almost undetectable after the beginning of the light period. To further determine if *TNY1* mRNA turnover was facilitated by light, RNA samples were collected under a shortened dark cycle, where cells were first synchronized under a standard 12hr:12hr light:dark regime, and then in the final cycle the length of the dark period was shortened to 3 hrs or 9 hrs, or lengthened to 15 hrs. When the light phase began early, *TNY1* mRNA disappearance was accelerated, while when the dark period was lengthened, *TNY1* mRNA persisted longer (Figure 4B, S4A). We conclude that some combination of light and/or cell growth promotes the reduction of *TNY1* mRNA through either accelerated mRNA turnover or decreased transcription.

The accumulation pattern of TNY1 protein throughout the cell cycle was determined in a wild-type culture synchronized under the standard 12hr:12hr light:dark diurnal cycle. Whole cell protein lysates were prepared from cells collected at different time points, fractionated by SDS-PAGE and probed for TNY1 protein on immunoblots that were also in parallel probed with histone H3 antibodies for cell number normalization. Two blots were prepared with different protein loading regimes on their respective gels. The first blot was made using equal numbers of cells loaded per lane to determine the absolute amount of TNY1 protein per cell (Figure 4C, S4B). During the G1 phase, cells increased in size by an average of eight-fold in size, but the absolute amount of TNY1 per cell remained relatively constant, and the signal only increased during S/M when mother cells began dividing. The other blot was from a gel loaded with equal amounts of total biomass per lane to determine the TNY1 concentration per cell (Figure 4D, S4C). Consistent with the first gel, the concentration of TNY1 per cell steadily dropped during G1 phase, as cells enlarged but no additional TNY1 was produced. In summary, cells are born with a fixed amount of TNY1 protein that is steadily diluted during G1 phase as cells grow, reaching its minimum concentration just prior to S/M during which its mRNA is transcribed and the protein is replenished in new daughters (Figure 4E).

We next determined how *TNY1* production scaled with mother cell size during S/M phase. Under the three regimes in Figure 4A, where mother cell size was increased or decreased based on time in light before division, the peak levels of *TNY1* mRNA increased or decreased compared with the standard condition in larger or smaller mother cells, respectively, suggesting that mother cell size or numbers of daughter nuclei may control *TNY1* mRNA production (Figure 4F, S4D, E). To determine if the subscaling of TNY1 is controlled by any feedback from size control regulators, we examined its levels in cell size mutants. TNY1 protein levels were determined in daughters produced from wild type, *mat3-4, dp1-1* and *cdkg1-2*. Interestingly, different sized daughter cells contained the same amount of TNY1 on a per cell basis (Figure 4G). Therefore, TNY1 production in each cell cycle is governed independently of the mitotic size control pathway, and its levels may instead be controlled by limiting factors that scale invariantly with cell size such as genomic template for *TNY1* transcription.

### TNY1 is limiting for size control

Size regulators that exhibit subscaling behavior are predicted to be dosage sensitive (23) (1, 6). To determine whether *TNY1* gene dosage might be limiting for size control, a set of isogenic diploid strains was constructed with genotypes *TNY1*/*TNY1, TNY1*/*tny1-1,* and *tny1-1*/*tny1-1* (Methods). Size profiles of daughters from synchronized diploid cultures of each strain were determined and compared to each other and to haploid strains. Daughters from the two homozygous strains were approximately twice the size of haploid daughters of the same genotype, while the heterozygous *TNY1*/*tny1-1* daughters were intermediate in size between the homozygous mutant and wild-type diploid strains (Figure 5A) and expressed less TNY1 protein than *TNY1*/*TNY1* diploids (Figure 5B). Further supporting the dosage sensitivity of *TNY1*, we found that among the meiotic progeny of *tny1-1*::*TNY1* (or *tny1-1::TNY1-HA*) rescued strains backcrossed to a wild-type parental strain, those that inherited both the wild type *TNY1* allele and the *TNY1* transgene were larger than those that inherited only the wild type *TNY1* allele or those that had the parental genotype of *tny1-1* with a *TNY1* rescuing construct (Figure S5A, B).

**Figure 5.**
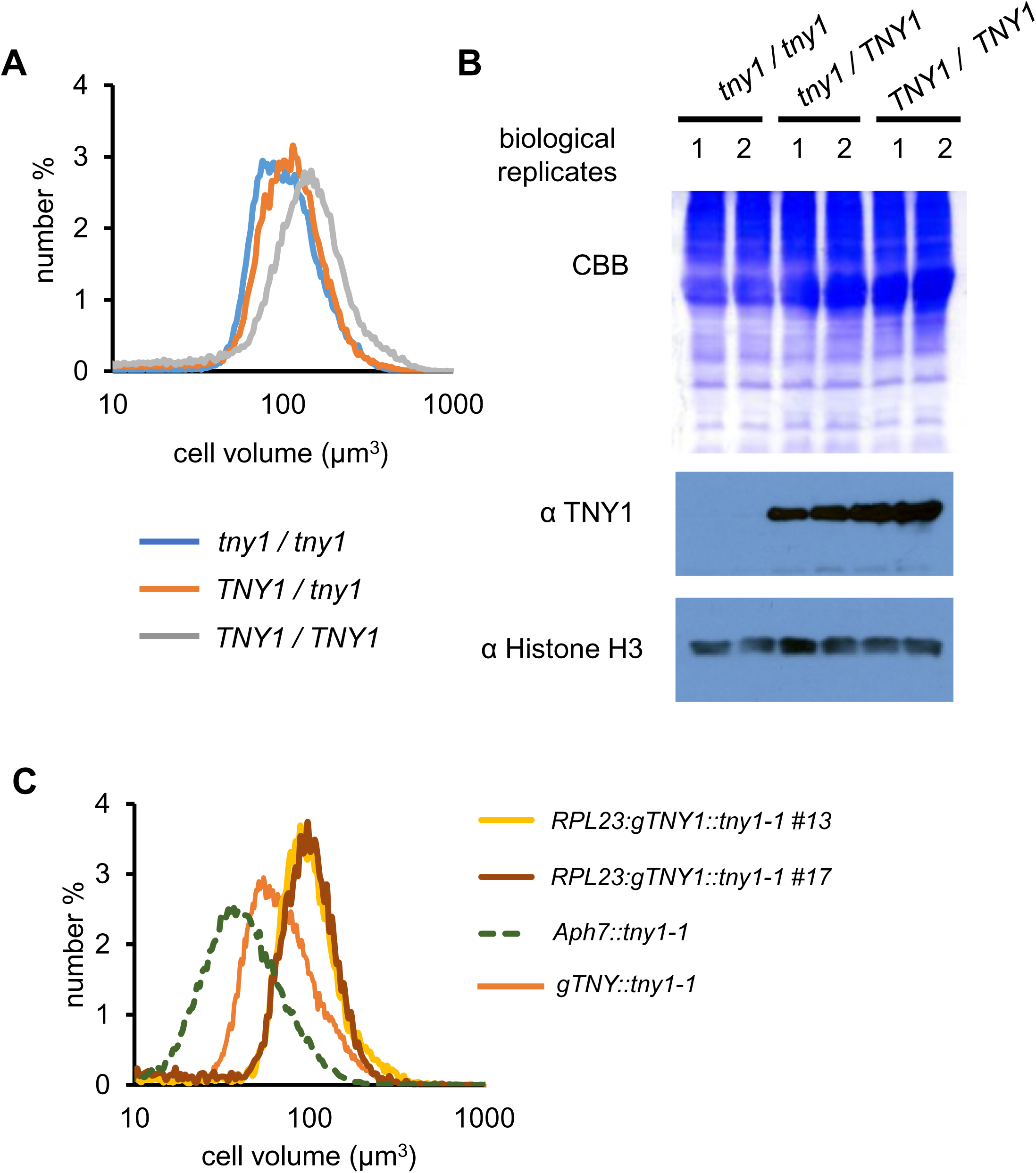
TNY1 is limiting in cell size control. (A) Size distributions of daughter cells of diploid strains with indicated genotypes. (B) Immunoblots and Coomassie gel were loaded and processed similar to those in Figure 4 (G) using two independently generated diploids (1 and 2) for each genotype. (C) Size distributions of synchronous daughter cells of two independent *RPL23:TNY1::tny1-1* rescued strains (#13, and #17), a control strain transformed with resistance marker only (*Aph7:tny1-1 #2*), and a *gTNY1::tny1-1* strain.

Besides altering gene dosage, we also generated a *TNY1* transgene driven by a previously characterized constitutive promoter/terminator from Chlamydomonas *RPL23* gene (24). This *RPL23:gTNY1:RPL23* construct was transformed into a *tny1-1* strain and transformants were tested for size phenotypes along with control transformants that received an empty vector with a selectable *aph7* marker conferring hygromycin resistance (25). Among independent *RPL23:gTNY1::tny1-1* transformants, ∼ 80% were rescued and were close to wild-type in mode size ∼ 80 µm^3^ (Figure S5C), while ∼ 20% showed a large-cell phenotype with a modal cell size > 100 µm^3^ that was never observed in rescue experiments using constructs with the controls (Figure 5C, Figure S5D). Large-sized *RPL23:gTNY::tny1-1* populations are always larger than the wild type strain throughout G1 (Figure S5E), while passing Commitment and entering S/M with similar timing as wild type and small-sized *tny1-1* mutants (Figure S5F). Taken together, these data indicate that dosage and expression level of TNY1 impact mitotic cell size control and are consistent with the subscaling behavior observed for TNY1 expression being an important contributor to size-dependent cell cycle control.

### TNY1 inhibits the accumulation of CDKG1 protein in post- mitotic cells

Previous work showed that mRNA and protein for the cell cycle activator CDKG1 are synthesized in a burst just before S/M phase and that CDKG1 protein is eliminated when cells exit S/M (2). Here we found that the negative regulator TNY1 functions genetically upstream of CDKG1 and that *TNY1* mRNA is upregulated during S/M, after *CDKG1* mRNA and protein are already present. These observations suggested a possible antagonistic relationship between TNY1 and CDKG1 in size control where TNY1 might limit production of CDKG1 or other cell cycle regulators during or after S/M phase. To test the effect of TNY1 on CDKG1 expression, wild-type and *tny1-1* strains were synchronized and *CDKG1* mRNA levels were measured in post-mitotic cells. Previously it was found that *CDKG1* mRNA and CDKG1 protein super-scale with mother cell size (2). In this experiment more *CDKG1* message was detected in *tny1-1* than in wild type, even though comparing the pre-division populations, *tny1-1* cells were smaller than wild type (Figure 6A). Since TNY1 is cytosolic, TNY1 is most likely to affect *CDKG1* mRNA levels by impacting message stability, though this finding does not rule out a possible role for TNY1 in translational control of CDKG1.

**Figure 6.**
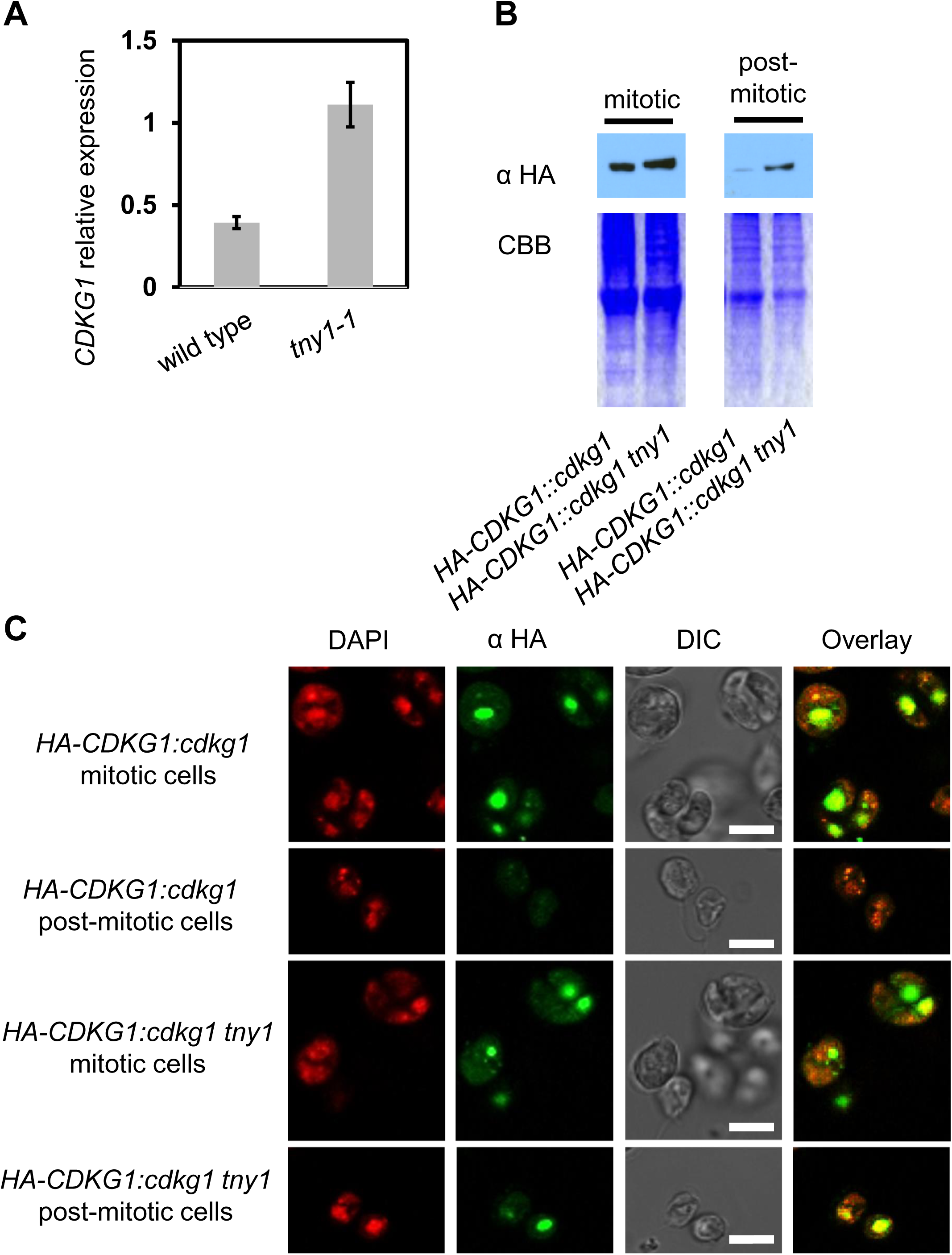
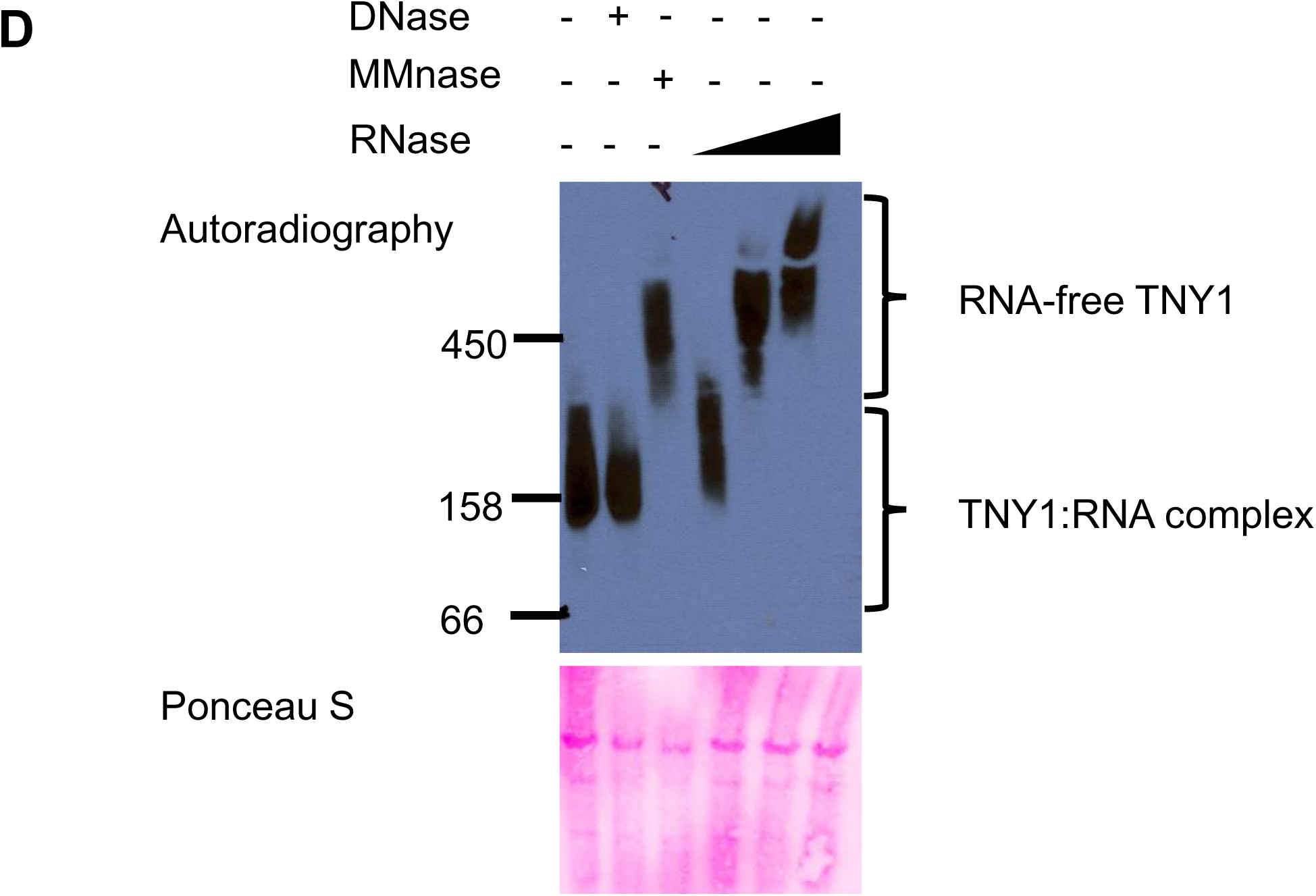
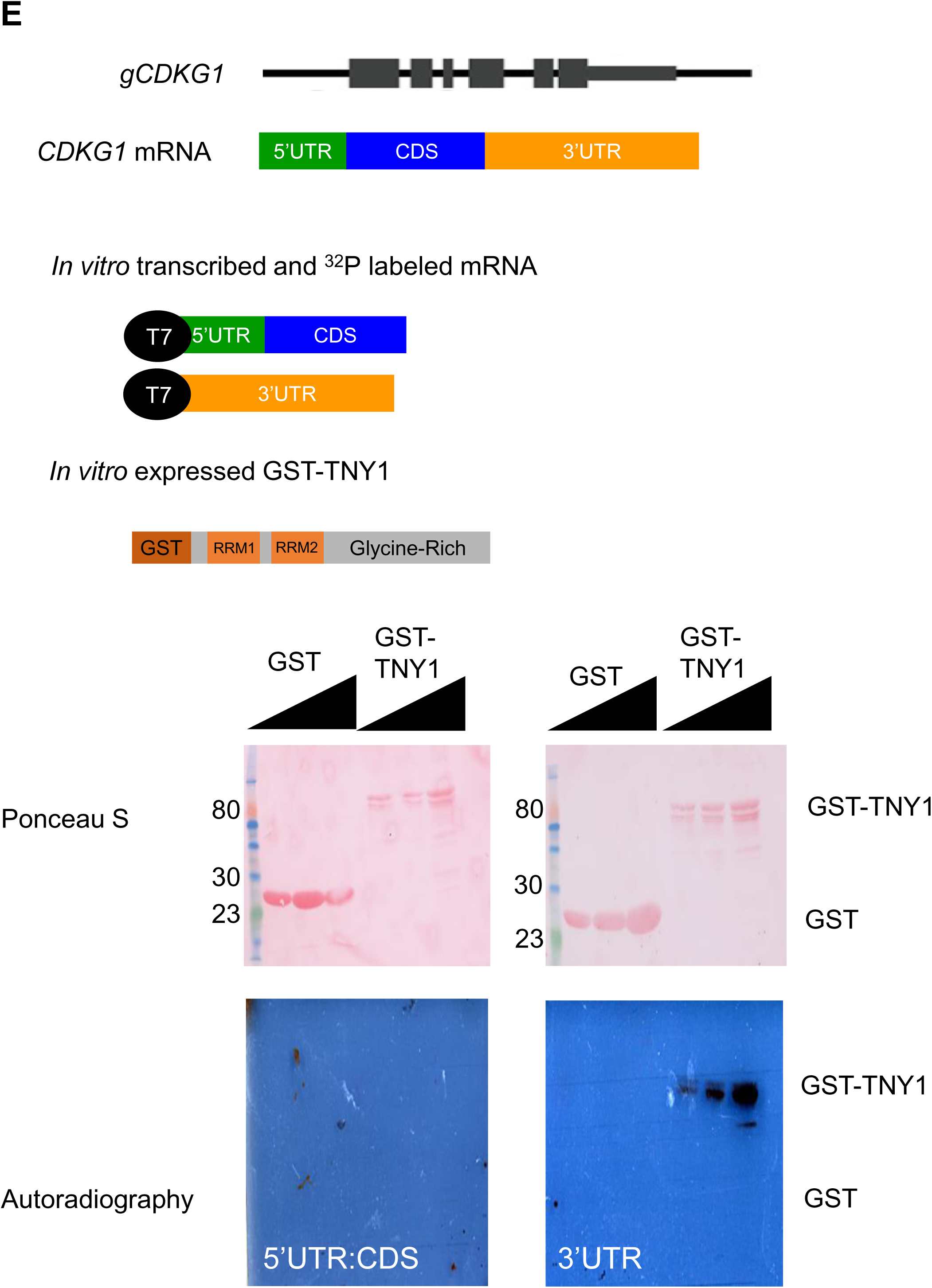
**TNY1 inhibits the accumulation of *CDKG1* mRNA and CDKG1 protein**. (A) qRT- PCR quantitation of average *CDKG1* mRNA level in daughter cells of wild type and *tny1-1* using two independently cultures (1 and 2) for each genotype. Error bars: high and low values of two biological replicates. The value of each biological replicate is calculated as the average of two technical replicates. (B) Immunoblots using synchronized strains of indicated genotypes loaded with equal numbers of cells per lane and probed with α-TNY1 (below) or stained with Coomassie blue above. Populations used are mitotic (ZT 13) or daughters using protein (ZT 24) lysate generated from synchronous mitotic and post-mitotic populations of indicated strains. Over accumulation of HA-CDKG1 was observed in the post-mitotic *HA-CDKG1::cdkg1 tny1* cells compared with a *cdkg1* rescue strain *HA-CDKG1::cdkg1*. (C) Brightfield and confocal immunofluorescence microscopy images of *HA-CDKG1::cdkg1* and *HA-CDKG1::cdkg1 tny1* cells. Synchronous mitotic and post-mitotic cells were fixed and immunostained for HA-CDKG1 (α-HA, pseudo-colored green), while the nuclei were stained with DAPI (pseudo-colored red). Note that some of the α-HA pixels were saturated, but all images were taken with similar settings and negative control experiments showed no nuclear staining. Scale bar = 10 μm.

To test the impact of TNY1 on CDKG1 protein abundance, *tny1-1* was crossed into a rescued *cdkg1* strain expressing an HA epitope tagged allele *HA-CDGK1* (2), so that CDKG1 protein levels could be assessed in a *tny1-1* strain background. CDKG1 content per cell was examined in protein samples from both mitotic cells and post-mitotic cells. *tny1-1* cells are smaller than wild type but showed higher levels of CDKG1 protein in mitotic and post-mitotic cells (Figure 6B). In addition, indirect immunofluorescence was used to detect HA-CDKG1 in mitotic and post-mitotic cells, where a clear HA-CDKG1 signal was present in nuclei of *tny1-1* daughters but not in the *TNY1* control strain (Figure 6C) (2). Together these data show that TNY1 limits the accumulation of both *CDKG1* mRNA and CDKG1 protein.

### TNY1 is part of an RNP complex and can bind to the 3’UTR of ***CDKG1* mRNA.**

The finding that cytosolic TNY1 could inhibit accumulation of nuclear-localized CDKG1 protein suggests a mechanism which might involve direct interaction of TNY1 with *CDKG1* mRNA. We first used native electrophoresis of whole cell extracts, and immunoblotting to determine if TNY1 might be part of a ribonucleoprotein complex (RNP). On native gels, TNY1 migrated near the 158 kDa marker which is slower than would be expected for free TNY1 (> 450 kDa). TNY1 migration was unchanged when extracts were pre-treated with DNAse, but when extracts were treated with microcoocal nuclease which digests both DNA and RNA, or by increasing amounts of RNAse, the TNY1 signal shifted to a slower moving complex migrating near the 450 kDa marker. These results suggest that TNY1 is associated with RNA *in vivo* as an RNP, and that the RNA component may contribute significantly to the negative charge state of the complex and impacting its migration rate on native gels (Figure 7A).

**Figure 7.**
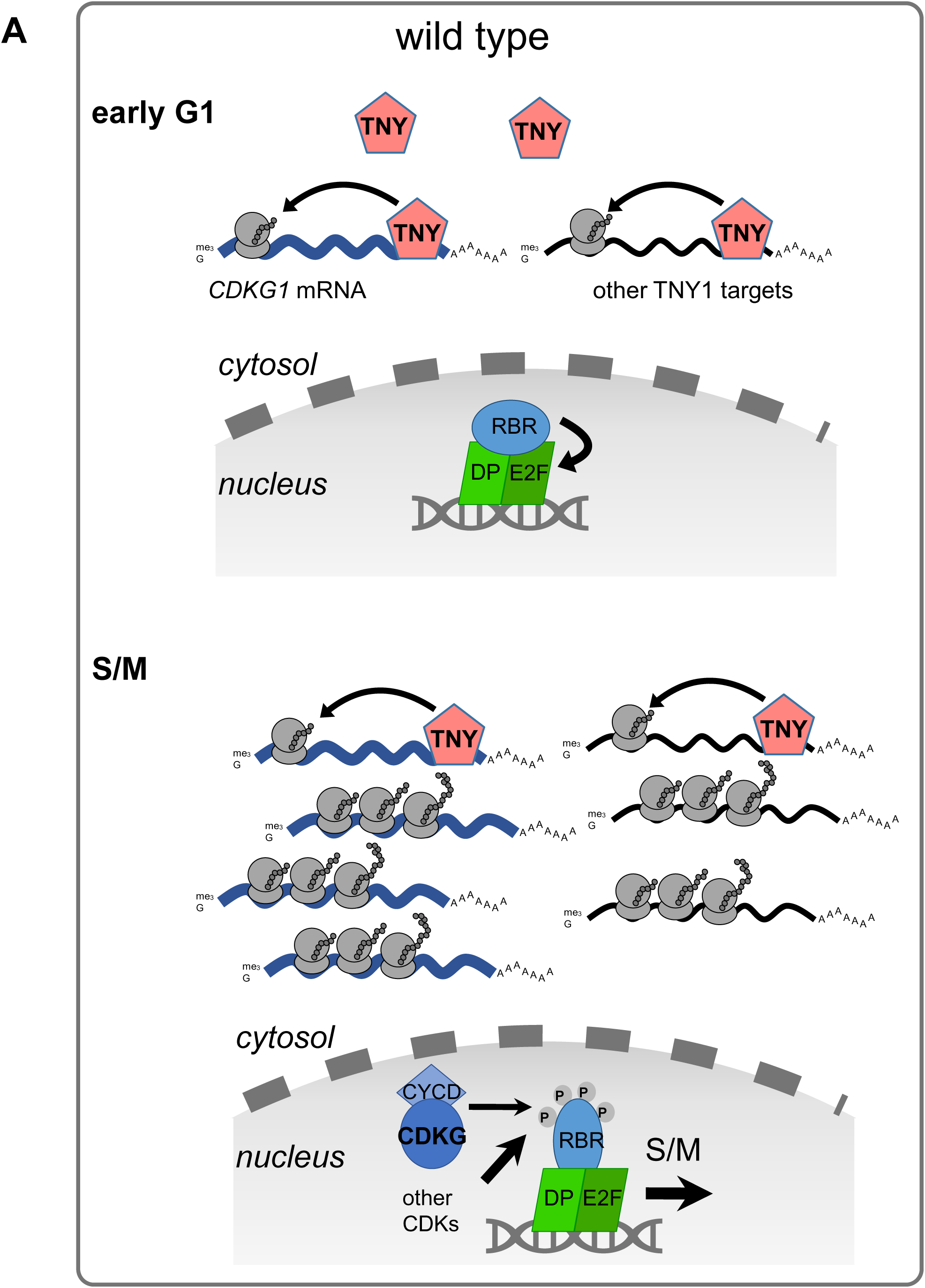
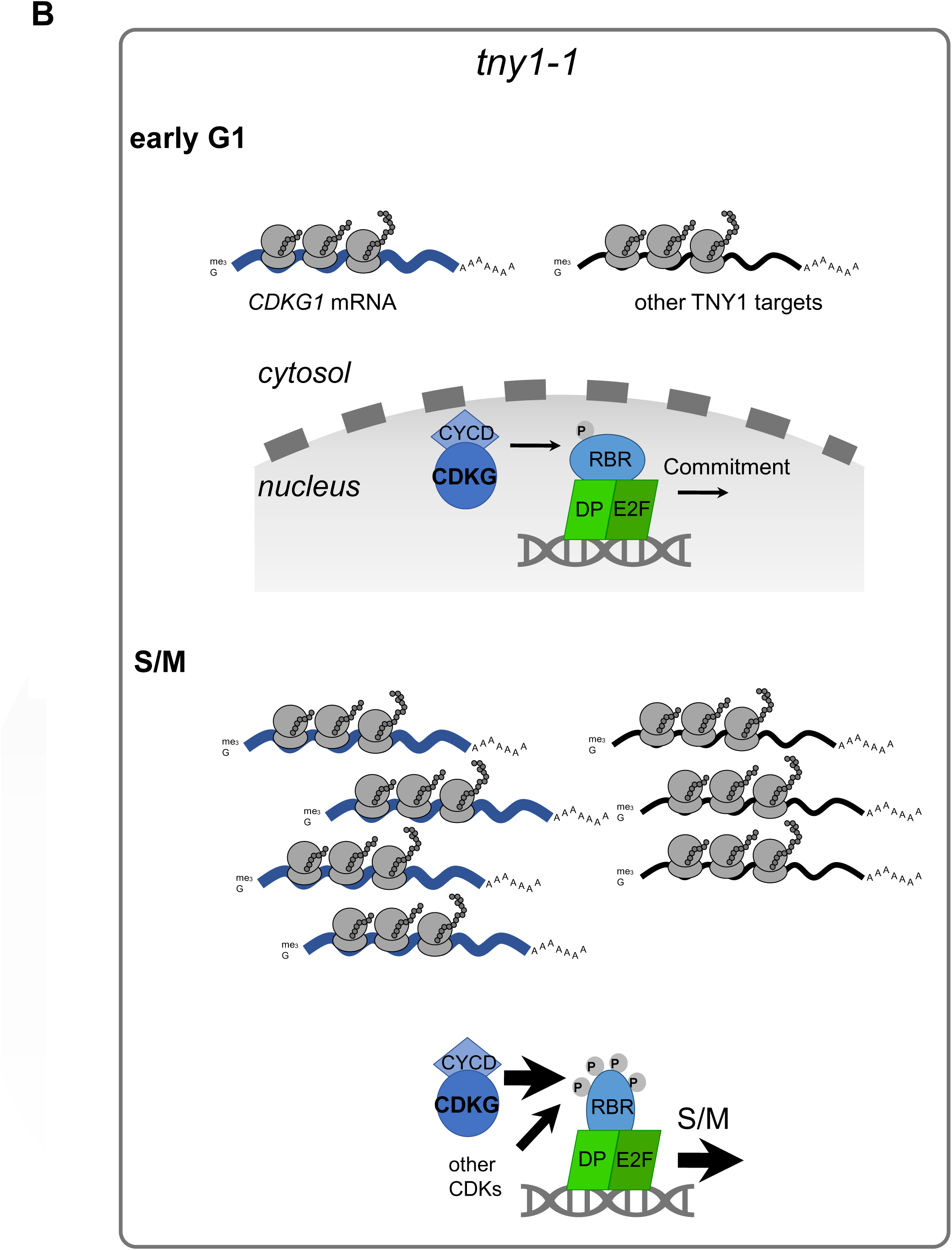
TNY1 binds to RNA including *CDKG1* 3’UTR. (A) TNY1 is part of a high molecular weight RNA-containing complex. Native gels were loaded with whole cell lysates of from a *gTNY1-HA::tny1-1* strain, fractionated and immunoblotted using α-HA. Lysates were pre-treated with different nucleases prior to loading as indicated above each lane with RNAse used at different concentrations indicated by the triangle with different RNAse concentration at 0.01 mg/mL, 0.1mg/mL, and 1mg/mL. The lower image is the same membrane stained with Ponceau S as a loading control. (B) GST-tagged TNY1 binds to ^32^P labeled *CDKG1* 3’UTR, but not to *CDKG1* 5’UTR and CDS in the northwestern assay. The total protein input was visualized by ponceau S staining.

A simple model for regulation of CDKG1 by TNY1 is direct binding of TNY1 to the *CDKG1* mRNA which has an unusually long (1.7kb) and uridine-rich (27% U versus 19% average for Chlamydomonas genes) 3’ UTR (26)— both rare features in Chlamydomonas mRNAs that tend to have shorter 3’ UTRs and overall GC-rich nucleotide composition. We attempted to detect TNY1 binding to *CDKG1* mRNA *in vivo* using RNA crosslinking and immunoprecipitation (RIP) (27) but were unable to amplify an enriched signal due to high background. Instead, we developed an *in vitro* assay where radiolabeled *CDKG1* mRNA fragments were used as a probe for binding to GST-TNY1 fusion protein or GST immobilized on a membrane (Methods) (28). Radiolabeled *CDKG1* mRNA was synthesized in two fragments, with the 5’ region including the 5’UTR and CDS in one fragment, and the 3’ UTR in a second fragment. After incubation of radiolabeled RNA with membrane-bound GST1-TNY1 or GST1 and washing, the signal was detected only for the 3’ UTR fragment binding to GST1-TNY1 (Figure 7B). These data indicated that TNY1 protein can bind RNA with sequence specificity, including sequences in the 3’ UTR of its likely target gene *CDKG1*.

## Discussion

In this study we identified a new Chlamydomonas sizer protein, TNY1, a hnRNP-related cytosolic RNA binding protein which functions as a negative regulator of cell size in a dosage-dependent manner. As with other size mutants in Chlamydomonas, *tny1-1* mutant cells retain relatively normal cell cycle progression kinetics, but do so with altered cell size setpoints for Commitment and for division number during S/M. *TNY1* mRNA and TNY1 protein are synthesized once per cell cycle during S/M phase, and TNY1 protein is at its highest concentration in newborn daughters. During G1 phase TNY1 absolute abundance remains constant, meaning that its cellular concentration drops as cells grow. This subscaling behavior appears to be important for size homeostasis since increased or decreased TNY1 dosage or expression impacts mitotic size control. The cell cycle activator and size regulator CDKG1, a D- cyclin dependent RBR kinase is a likely direct target of TNY1 repression since ectopic accumulation of CDKG1 protein and mRNA was observed in *tny1-1* mutants, and TNY1 protein could interact specifically with the 3’UTR of the *CDKG1* mRNA, possibly as a translational repressor or destabilizing factor.

Together these data suggest a model where TNY1 controls cell division by modulating the accumulation of a limiting activator protein, CDKG1, and possibly other limiting cell cycle regulators (Figure 8). This modulation might occur in at least two ways. During G1 phase, CDKG1 is not detectable and does not seem to play a normal role in cells passing Commitment, but in a *tny1-1* mutant its inappropriate expression in G1 phase could change the Commitment threshold size by contributing to the premature inactivation of the RBC which controls Commitment cell size (15, 16). Just prior to S/M phase, the absence of TNY1 may cause the production of extra CDKG1 leading to increased division number during S/M, or it may cause extra divisions by preventing the timely removal of CDKG1 which normally accompanies mitotic exit (Figure 8B). *In vivo* binding studies to determine the timing of when TNY1 associates with *CDGK1* mRNA, and to identify other direct RNA targets of TNY1 will be useful for testing the direct repression model for cell size control.

**Figure 8.** Model for subscaled TNY1 as a regulator of size-dependent cell cycle progression. (A) In wild-type cells during early G1 phase (top half) cytosolic TNY1 binds the 3’UTR of *CDKG1* mRNA and possibly other targets and prevents premature expression. Prior to and during early S/M phase (bottom half) *CDKG1* mRNA and other target mRNAs outnumber TNY1 protein which is at its lowest concentration. Translation of CDKG1 drives size-dependent cell cycle progression through phosphorylation of RBR by CDKG1/D-type cyclins and other mitotic kinases in the nucleus (2). (B) In *tny1* mutants some CDKG1 is inappropriately produced in early G1 phase (top half) and may prematurely push cells to Commitment at a smaller size through ectopic phosphorylation of RBR. During S/M phase (bottom half) the absence of TNY1 allows extra CDKG1 to accumulate causing an imbalance in size sensing and more cell divisions than in equivalent-sized wild-type mother cells.

Evidence for cell size checkpoints based on some form of protein subscaling has been found in different eukaryotic taxa, including fungi, animal cells and plant meristems (Figure S6) (7, 8, 10, 29). An appealing property of subscaled proteins is that their absolute abundance in a cell can act as a denominator for perceiving changes in cell size whose proxy is a protein or other molecule whose cytoplasmic concentration is constant. Interestingly, in the above examples subscaling could be directly tied to nuclear function via DNA or chromatin (6). In budding yeast, nuclear Whi5 protein binds to and inhibits the DNA bound transcription factor SBF, a key activator of S phase transcription. While some regulation of Whi5 abundance may occur based on synthesis of Whi5, it is also limited by chromatin binding (8, 29, 30). Similar findings were made for the RB protein in mammalian cells which is a functional analog of Whi5 for S phase transcription (10). In plants, chromatin binding by the CDK inhibitor KRP4 coupled with elimination of excess unbound KRP4 allows daughter cells to be apportioned with a fixed amount of KRP4 that acts as a concentration dependent inhibitor of the cell cycle in the subsequent G1 phase and ensures that S phase entry occurs at a constant average cell size regardless of daughter cell sizes (7). Here we found that subscaling can also occur for a cytosolic protein, TNY1, that has no direct connection to the nucleus or chromatin. This finding raises the question of how TNY1 synthesis is controlled and how its levels can be modulated so that daughters always contain the same amount of TNY1. One way to achieve a fixed dose of TNY1 per cell would be if production of *TNY1* mRNA is limited by *TNY1* gene copy number in daughters and not influenced by cell size related factors (e.g. transcription factor abundance, co-activator abundance) (23), but this remains to be determined. Supporting this idea, TNY1 absolute abundance in daughters was not influenced by cell size mutants that caused production of large or small daughters. To date, TNY1 is the only cell cycle regulatory protein in Chlamydomonas known to subscale. The RB complex is downstream of TNY1 in Chlamydomonas, but MAT3/RBR increases in abundance during G1 phase (2, 16) and does not show dosage sensitivity for size control as its mammalian homolog RB and its yeast counterpart Whi5 do. Thus, the systems-level target for subscaling of size control is not conserved between algae and these two members of the opsithokont phylum.

Interestingly, TNY1 shares some similarity to budding yeast Whi3, an RNA binding protein and negative cell cycle regulator that functions in part by restricting expression of the limiting G1 cyclin Cln3 (31, 32). In budding yeast, Whi3 represses the function of Cdc28-Cln3 by retaining Cdc28-Cln3 complexes in the cytoplasm in G1 phase (33). Whi3 does not impact the abundance of Cdc28 but does represses *CLN3* mRNA abundance and translational efficiency (34). In Chlamydomonas, TNY1 functions upstream of CDKG1 and appears to repress the accumulation of *CDKG1* mRNA and CDKG1 protein. Unlike Whi3, cytosolic TNY1 does not impact the nuclear localization of CDKG1. Musashi proteins (MSIs) are metazoan hnRNP that play a role in stem cell maintenance and proliferation (35). While the targets of MSIs are not fully defined, they primarily bind to 3’ UTRs of mRNAs and regulate mRNA stability and/or translation (36, 37). Future work aimed at systems-level understanding of cell size regulatory networks may reveal additional parallels for RNA binding proteins such as TNY1 in cell size and cell cycle control.

## Methods

### Chlamydomonas strains and growth conditions

Strains were maintained on Tris-acetate-phosphate (TAP) + 1.5% agar plates. For synchronous growth they were cultured at 25°C in Sueoka’s High-Salt-Media (HSM) liquid media (38) with diurnal cycles as indicated and 300 µE total light intensity (50% Blue:50% Red = 150 µE blue at 465 nm and 150 µE red at 625 nm LED lights) bubbling with 1% CO_2_. Diurnal light regimes used are described in figure legends and text.

Gamete generation, mating, and zygote germination were performed following standard protocols (39–41). Segregation analysis was done with randomly selected progeny from mating. Dark-shift experiments, commitment assays, and cell-size distribution measurements with a Coulter Counter (Beckman Multisizer 3) were conducted as described previously (42). The mode size for a particular cell population was determined from the peak of a smoothed log-normal size histogram curve. Modal cell size and mean cell size were calculated manually using cells within the size range 20 µm^3^ - 2000 µm^3^. Particle sizes above and below this range are rare, and mostly consist of small debris or large clumps.

### Chlamydomonas transformation

Cells were cultured asynchronously at 25°C in TAP liquid media (43) with constant light 100 uE total light intensity (50% Blue:50% Red – 50 µE blue at 465 nm and 50 µE red at 625 nm LED lights) bubbling with filtered air. Cells were transformed using electroporation as previously described (24). Transformants were plated on TAP agar plates with either 15 µg/mL of paromomycin or 25 µg/mL of hygromycin depending on selection markers.

### Forward genetic screen for size mutant and mapping of *tny1- 1*

Wild-type strain CC124 was subject to an insertional mutagenesis using vector pSI103 (25) linearized with NotI and transformed using the glass bead method (44) with selection on TAP agar plates containing 12 µg/mL paromomycin. Transformants were picked and re-grown in individual wells of 96 well plates, then stamped onto TAP agar plates using a 48 Multi-Blot Replicator (, which delivers ∼3 μL hanging drop) on a light shelf at 25°C for 6 days.

Approximately 1/3 of each stamped spot was removed with a toothpick and resuspended in nitrogen-free HSM in a new 96 well plate to create a gamete suspension. Gametes were then checked for cell size using a Coulter Counter. Confirmed mutants were then crossed to wild-type strain CC125, and progeny were tested for linkage of the suppressor phenotype to the pSI103 insertion. The *tny1-1* insertion site was determined by sequencing junction fragments from ligation mediated PCR (45), and the insertion site was confirmed using genotyping primers for *tny1-1* (Supplementary Table).

### Diploid generation

Diploid selection was done by plating crosses (described below) shortly after mating on double selection plates with antibiotics to select for both parent stains. Wild type *TNY1 TNY1* vegetative diploids were generated by a mating between wild type CC-1039 wild type (Sager’s 21 gr) (*NIT1 NIT2* (Nit+) *MT+*) and wild type CC124 transformed with pKS-aph7’’-lox (46) (*MT-*, hygromycin resistant, *nit1 nit2* (Nit-)) with selection on 25 µg/mL hygromycin and nitrate as the only nitrogen source. Heterozygous *TNY1 tny1* vegetative diploids were generated by a mating between wild type CC-1039 and *tny1-1* (*MT-*, paromomycin resistant) with selection on paromomycin with nitrate as the only nitrogen source. Homozygous *tny1 tny1* vegetative diploids were generated by mating between a *tny1 MT+* Nit+ segregant from a cross with CC-1039 and *tny1* transformed with Aph7, with selection on plates with 25 µg/mL hygromycin and nitrate as the only nitrogen source. Diploid candidates validated by genotyping with mating-type loci primers (47).

### Rescue of *tny1-1*

A 3.4 kb fragment containing the full-length genomic region of *TNY1* was amplified from genomic DNA using primers TNY KpnI/TNY NdeI listed in Supplementary Table. The amplified fragment was digested with KpnI/NdeI and ligated into KpnI/NdeI digested vector pHyg3 linearized with the same restriction enzymes to generate *tny1* rescue construct *pTNY1*. A triple hemagglutinin epitope tag (3xHA) was inserted into pTNY1 just before the stop codon into a BgllI site created by overlapping PCR with 2 fragments amplified with oligos TNYKpnI/ TNYBglIIRev and TNY BglIIF/TnyNdeIIF (Supplementary Table). A triple HA epitope tag (3xHA) was amplified from 3xHA-MAT3 (16) with oligos HABglII-F/HABglIIR cut with BglII and inserted at the BglII site just before the translation stop codon.

*pTNY1* or *pTNY1-3xHA* were transformed into *tny1* by electroporation as described above with selection on TAP agar with 30 μg/mL hygromycin. Individual transformants were picked into 96 well plates and screened for gamete cell sizes as described above for screening insertional mutants. TNY1 expression was confirmed by immunoblotting (see below).

### Mis-expression of *TNY1*

To generate pRPL23-TNY1, full genomic *TNY1* fragment between the start and stop codons was amplified with primers BamHI TNY1 F and Xho1 TNY1 R (Supplementary Table) from *tny1* rescue construct pTNY1. The amplified *TNY1* fragment was digested with BamH1 and Xho1 and inserted into pRPL23:Luc:RPL23 (24), then recombined with plasmid pKS-aph7’’-lox (46) to generate pRPL23-TNY1-aph7. pRPL23-TNY1-aph7 or pKS-aph7’’-lox (negative control) were transformed into *tny1-1* by electroporation (see above). Transformants were selected on TAP agar plates containing 25 µg/mL hygromycin.

### Phylogenetic analysis of TNY1 and hnRNP proteins

BLAST searching was done within NCBI or on Phytozome (17) using Chlamydomonas TNY1 protein sequence as a query to find high-scoring hits in plants, green algae and holozoans.

Tandem RNA binding domain proteins are found in most eukaryotes, with several representatives besides TNY1 within Chlamydomonas. However, the top BLAST hits for TNY1 were found outside of Chlamydomonas as single best hits within other species of green algae, including three representative volvocine algal species (*Gonium pectorale*, *Tetrabaena socialis*, *Volvox carteri*). The sequences were aligned using MAFFT within Guidance2 (48), and the well-supported portion of the alignment of 158 residues containing the RNA binding domains was retained for phylogenetic analysis. Some duplicates and very closely related sequences were removed to reduce redundancy, with a final group of 39 proteins used for phylogenetic reconstruction. Evolutionary models were tested using Modeltest-NG (49), with the best model being LG+G(1.46)+I(0.08). A maximum likelihood phylogeny was estimated using W-IQ-tree (50) with approximate likelihood ratio testing of branch support.

### TNY1 antibody generation

A full length *TNY1* cDNA was amplified with primers TNY1-1F and TNY1-1R (Supplementary Table) from cDNA prepared using RNA from wild-type strain CC124 and inserted into pGEM-T easy vector (Promega) to generate pGEM-TNY1. After verification by Sanger sequencing the *TNY1* cDNA fragment was released by digestion with NdeI and XhoI (NEB) and inserted into vector pET28a digested with NdeI and XhoI. The construct was transformed into *E.coli* strain BL21 codon plus-RIL (DE3) (Agilent technologies). Induction of recombinant TNY1 expression in *E. coli* and purification of insoluble 6xHis-TNY1 was performed under denaturing conditions as described previously (16). Purified 6xHis-TNY1 was cut out from a Coomassie blue stained SDS-PAGE gel and sent to Cocalico Biological Inc. to generate rabbit polyclonal anti-sera. Polyclonal antibodies were affinity purified with AminoLink Plus Resin (Thermo Fisher) coupled to purified GST-TNY1 (see below).

### Protein extraction and western blotting

Chlamydomonas cultures were grown as described above and harvested by centrifugation at 4000g for 5 min after adding tween-20 to a final concentration of 0.005%. Pellets were washed in PBS and resuspended in lysis solution (1xPBS pH 7.4, 1x Sigma plant protease Inhibitor, 5 mM Na_3_VO_4_, 1 mM NaF, 1mM Benzamidine, 500 mM PMSF, 1 µM ALLN, 1 µM MG-132) to a final concentration of 5x10^8^ cells/mL, and immediately frozen in liquid nitrogen. Pellets were thawed on ice and centrifuged at 12,000 g for 10 min at 4°C. Supernatants were transferred into a new tube and boiled for 5 min in SDS protein loading buffer as whole cell lysates. Total protein was separated on 12% SDS-PAGE gels and wet-transferred to PVDF membranes at 50 Volt for 1hr. Membranes were blocked in PBS containing 9% nonfat dry milk, incubated overnight with primary antibodies (1:5000 α-TNY1, 1:10,000 Roche Rat-anti-HA, high affinity 3F10, or 1:50,000 Invitrogen α-Histone H3) in 5% non-fat dry milk. Membranes were then washed in PBS containing 0.1% Tween 4 x 15 min, incubated at room temperature with secondary antibodies coupled to horseradish peroxidase (1:20,000 Thermo Fisher goat-anti-rabbit, or 1:20,000 Millipore Sigma goat-anti-rat in 5% nonfat dry milk). Membranes were washed again in PBS containing 0.1% Tween 4 x 15 min, then subject to chemiluminescent detection using autoradiographic film or a Bio-Rad quantitative imaging system (Chemi DocTM XRS+ Imaging System).

### qRT-PCR

Total RNA samples were extracted at different time points from synchronized strains using a Trizol-like reagent following the method of (15) then digested with RNase-free Turbo DNase following the manufacturer’s protocol. 4 µg total RNA was reverse transcribed with oligo dT and random hexamers (9:1) using Thermo Script Reverse Transcriptase at 25°C for 10 min, 42°C for 10 min, 50°C for 20 min, 55°C for 20 min, 60°C for 20 min, 85°C for 5 min. SYBR-Green based qPCR reactions in two technical duplicates of two biological replicates were performed and quantitated in a Bio-Rad CFX96 system. Each 10 μL reaction contained 0.1 µL cDNA, 1x Invitrogen Taq buffer, 3.5 mM MgCl_2_, 0.5x SYBR Green I, 0.05% Tween 20, 0.05 mg/mL BSA, 5% DMSO, 200 µM dNTPs, 0.3 µM primers, and 5U of Invitrogen Taq DNA polymerase.

Expression was normalized against *GBLP* (GenBank NC_057009.1) as an internal control. The melting curve was examined for each reaction to ensure that no primer dimers or non-specific PCR products were present. qPCR primers for *CDKG1*, *TNY1,* and *GBLP* can be found in the Supplementary Table.

### Light microscopy

Chlamydomonas cells were fixed in 0.2% glutaraldehyde final concentration. Cells were mounted on slides and imaged with a Leica DMI 6000 B microscope with a 63* oil objective and DIC optics with images taken using a Photometrics Coolsnap HQ2 CCD camera.

### Immunofluorescence microscopy

*TNY1-HA::tny1, HA-gCDKG1:: cdkg1-2* (2), or *HA-gCDKG1:: cdkg1-2 tny1-1* strains were synchronized as described above on a 14hr light: 10hr dark diurnal cycle. S/M phase cells were collected at ZT 15 hrs and daughter cells at ZT 23 hrs. Cells were centrifuged and collected in an Eppendorf tube, fixed with 2% paraformaldehyde in PBSP (1x PBS pH7.4, 1 mM DTT, 1x Sigma plant protease inhibitor cocktail) for 30 min on ice. Fixed cells were extracted in cold methanol 3 x 10 min at -20°C and rehydrated in PBSP for 30 min on ice. Cells were blocked for 30 min in blocking solution I (5% BSA and 1% cold water fish gelatin in PBSP) and 30 min in blocking solution II (10% goat serum, 90% blocking solution I). Cells were incubated overnight with primary antibody α HA antibodies (Roche 3F10) (1:1000 dilution in 20% blocking solution I) at 4°C, then washed 3 x 10 min in 1% blocking solution I at room temperature. Cells were then incubated with 1:1000 Alexa Fluor 568 conjugated goat anti-mouse IgG in 20% blocking solution I for 1 hr at 4°C and then incubated with 4’,6-Diamidino-2-Phenylindole, Dihydrochloride (DAPI) at a final concentration of 5ug/mL for 5 min. Cells were washed in 1 x PBS for 3 x 10 min. Cells were mounted in 9:1 Mowiol: 0.1% 1, 4-phenylenediamine (PPD), and imaged with a Leica DMI 6000 B microscope with a 63x oil objective (NA 1.40) and a Photometrics Coolsnap HQ2 CCD camera. Fluorescence illumination was provided by a metal halide lamp (Prior Lumen 200 Fluorescence Illumination Systems) using a Leica A4 filter cube (ex 360/40; em 470/40) for DAPI imaging and TX2 filter cube (ex 560/40; em 630/75) for detection of TNY1.

### Construction of a TNY1-mCherry expressing strain

To generate a fluorescence protein-tagged *tny1* complemented strain, a *pTNY1:gTNY1-GFP- TNY1 3’ UTR* construct was generated first. Chlamydomonas codon optimized GFP fragment (SpeI-SacI-BamHI-GFP-Xba-Xho-EcoR-NcoI) was amplified from *pMF124cGFP* (51) and digested by SpeI and NcoI, followed by insertion into *RPL23:Luc:RPL23* which is digested by XbaI and NcoI. A fragment of *pTNY1:gTNY1,* including the promoter region, 5’UTR, and exons and intron of genomic TNY1, was amplified and digested with SacI and BamHI, and inserted into the above modified GFP plasmid. TNY1 3’UTR and terminator region was amplified and digested with XbaI and EcoRI, followed by insertion into the above *pTNY1:gTNY1-GFP* backbone.

Chlamydomonas codon-optimized mCherry was amplified using a primer set of BamH1 mCherry F and XbaI mCherry R (Supplementary Table) from pLM006 (20), digested with BamH1 and Xba1, then used to replace GFP in the plasmid *pTNY-GFP* digested with BamHI and XbaI to create plasmid *pTNY1-mCherry. pTNY1-mCherry* was transformed into *tny1-1* and rescued transformants were identified by measuring gamete sizes as described above and then confirmed by immunoblotting and measuring sizes of vegetative daughter cells. for *TNY1- mCherry* using α TNY1 antibody.

### Live cell confocal fluorescence microscopy

*pTNY1-mCherry* expressing transformants or a rescued negative control strain expressing *pTNY1-HA* were synchronized and harvested throughout the multiple fission cell cycle. Live cells were immobilized on a very thin layer of TAP agar on a glass slide, and topped with a coverslip, which was sealed with PicoDent following the manufacturer’s instructions (https://www.picodent.de/). Cells were imaged using a Leica SP8-X confocal microscope equipped with a white light laser and a 405 nm diode laser. TNY1-mCherry was detected with 570 nm excitation and a 550-650 nm emission window. Fluorescence lifetime gating 0 - 4.9 ns was used to remove most of the chlorophyll background signal. Chlorophyll was detected using 405 nm excitation and a 676-704 nm emission window.

### Native gel separation and detection of TNY1 RNP complexes

50 mL samples from Chlamydomonas cultures at 10^6^ cells/mL were mixed with Tween-20 to a final concentration of 0.005% and collected by centrifugation at 4000 g for 5 min. Pellets were washed in PBS and resuspended in lysis solution (1xPBS pH 7.4, 1x Roche plant protease Inhibitor, 1 mM PMSF) to a final concentration of 5x10^8^ cells/mL, and immediately frozen in liquid nitrogen. Pellets were thawed on ice and centrifuged at 12,000 g for 10 min at 4°C.

For RNA binding assays, 20 μL of supernatant was incubated with different RNAse dilutions 1:10, 1:100 or 1:1000 (stock 10 mg/mL, NEB) or with 1:10 Dnase I (stock 2 U/μL, Roche), and micrococcal nuclease (stock 2000 U/μL, NEB). 6* protein loading sample buffer without DTT or SDS was added to samples before loading into a precast native 4-12% tris glycine gel (Invitrogen) without SDS in Tris-Glycine running buffer. A mixture containing aldolase, BSA and ferritin was used as a molecular weight marker. Native PAGE gels were transferred to nitrocellulose membranes in 25 mM Tris, 192 mM glycine, 20% methanol. Blots were blocked in 1x PBS with 5% non-fat dry milk for 1h at room temperature and incubated with 1:2500 anti- TNY diluted in PBST (PBS + 0.05% Tween-20) with 3% dry milk at 4°C overnight. After washing in PBST for 3* 10 min, the blot was incubated with horseradish peroxidase (HRP) conjugated goat-anti-rabbit-IgG (1:5000, Pierce ECL) for 1hr at RT, then washed in PBST for 3* 10 min, and processed for chemi-luminescence (Luminata forte, Millipore).

### 32P RNA radio-labeling

*CDKG1* DNA for *in vitro* transcription was amplified from genomic DNA with oligos containing a T7 promoter (Supplementary Table). ^32^P labeled RNA was generated/ transcribed *in vitro* using a Maxiscript kit in the presence of α-^32^P-CTP (NEN) according to manufacturer instructions.

Each 25 μL reaction had the following components: DNA template 0.5ug, 10x Transcription buffer 2 μL, 0.5 mM ATP, 10mM GTP 1 μL, 10mM UTP 1 μL, 500uM CTP 1 μL, ^32^P-CTP 2 μL (10 mCi/mL), 2 μL T7 RNA polymerase. After 1hr reaction at 30°C, the mixture was treated with DNAseI (ambion) and purified with Sigma post reaction clean-up columns SigmaSpin™ to remove unincorporated nucleotides. RNA integrity was visualized by separating a sample of the RNA on a urea denaturing 4% polyacrylamide gel followed by autoradiography.

### GST-TNY recombinant protein expression

The *TNY1* cDNA coding sequences were cloned into the Gateway pDEST15-GST (glutathione S-transferase) plasmid using the procedures recommended by the manufacturer (Invitrogen) with oligos listed in Supplementary Table 1. Mutations in TNY1 RRM1 and RRM2 motifs, RRM1* and RRM2* respectively, were introduced with Quick change mutagenesis kit with oligos listed in Supplementary Table. To generate TNY1 RRM1* RRM2* double mutant, TNY-RRM1* was digested with Nde1/Sal1 and ligated into TNY RRM2* in pENTR, and then moved to Gateway pDEST15 destination vector. GST-TNY constructs were transformed into *E.coli* BL21 codon plus-RIL strain (Agilent Technologies). Cells were grown in LB media and induced for 5 hrs at 30°C with 0.5 mM isopropyl-β-d-thiogalactopyranoside (IPTG) when cultures reached an O.D.600 of 0.5. After induction, cells were harvested by centrifugation and dry cell pellets stored at -80 °C. Frozen cells were thawed on ice and resuspended in one-tenth original culture volume of EB (100 mM Tris-HCl, pH 8.0, 500 mM NaCl, and 10 mM imidazole), sonicated eight times for 2 min each on ice with a Branson sonicator (50% power with a duty cycle of 0.5s on and 0.5 s off) followed by supernatant clearance by centrifugation at 12,000g for 10 min. GST- TNY recombinant proteins were purified from the soluble fraction using Glutathione Sepharose beads (Amersham) following the product manual.

### TNY1 RNA binding assay

Equal amounts of GST purified proteins estimated based on Ponceau S staining were separated by SDS-10% PAGE and transferred to a nitrocellulose membrane (0.22-m pore size) and stained with Ponceau S. The membrane was incubated at 4°C overnight with re-naturation buffer: 50 mM tris-HCl pH 7.5, 100 mM KCl, 1% Triton X-100 and 10% glycerol. After re- naturation, the membrane was incubated for 1hr with reactivation buffer (Tris-HCl pH 7.5, 0.1 % triton X-100, 10% glycerol) at room temperature, blocked for one hour with yeast tRNA (80 μg/mL) in reactivation buffer followed by incubation with ^32^P labeled RNA in reactivation buffer for 3 hrs. Membranes were washed 4 times with reactivation buffer and exposed to X-ray film for 2 days at -80°C before development.

## Acknowledgement

We thank Tuya Wulan, Fuqin Sun, Richard Davenport, Thomas Connell, Kerri Husa, Nazifa Hoque, Jie Li, Dylan Wetzel, Hunter Draffen, Brooke Harris, Zach Jaudes, and Chris Reynolds for laboratory support. We thank Dr. Kirk Czymmek, Dr. Anastasiya Klebanovych, and Dr. Howard Berg at the Danforth Plant Science Center for the guidance on microscopy. This work was funded by NIH R01GM092744, NIH R01GM126557, NSF MCB 1515220 to Dr. James Umen, and facilitated by the research instruments acquired through NSF DBI 2018962 to the Danforth Plant Science Center.

## Supporting information

**Figure S1.**
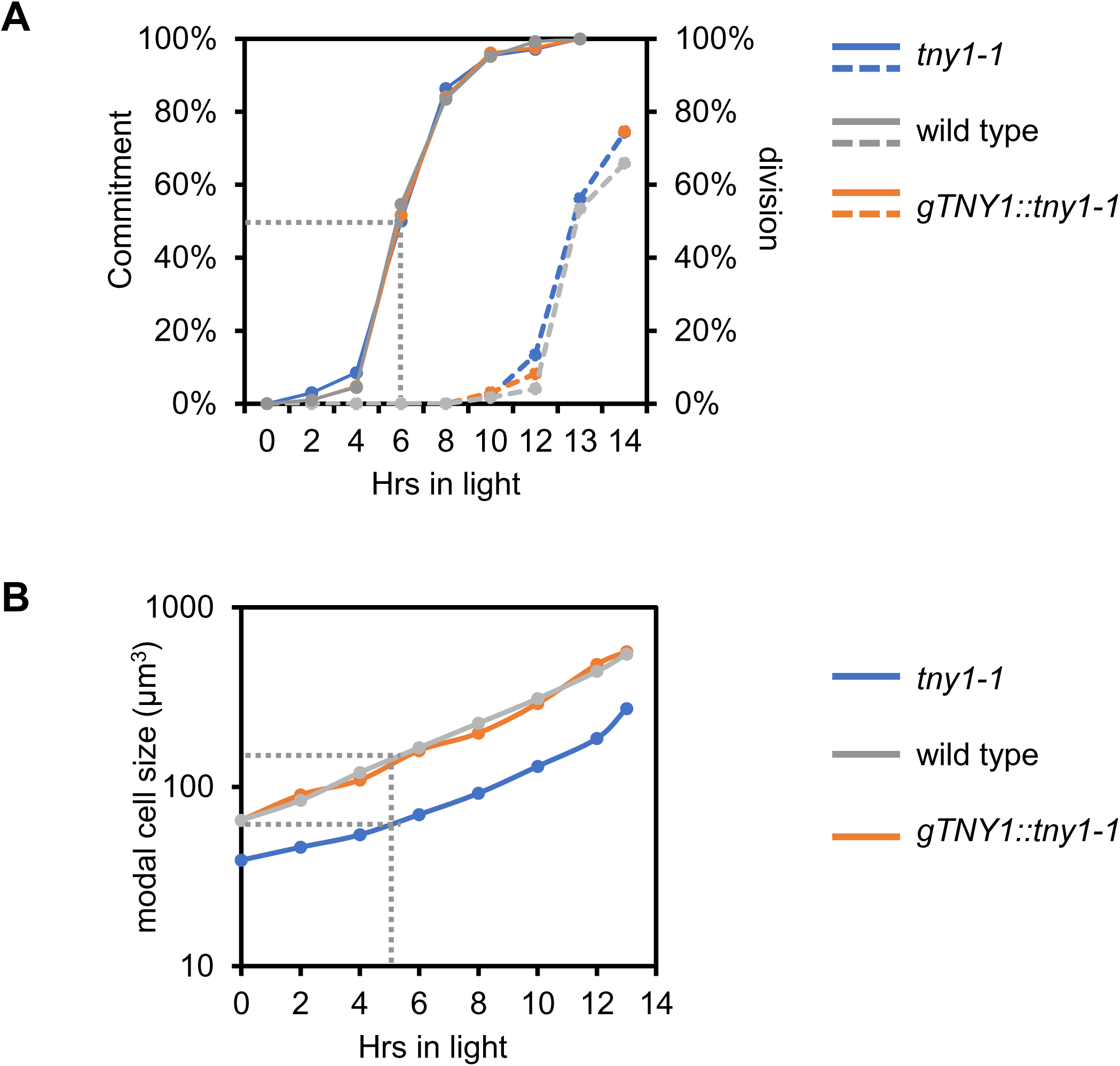
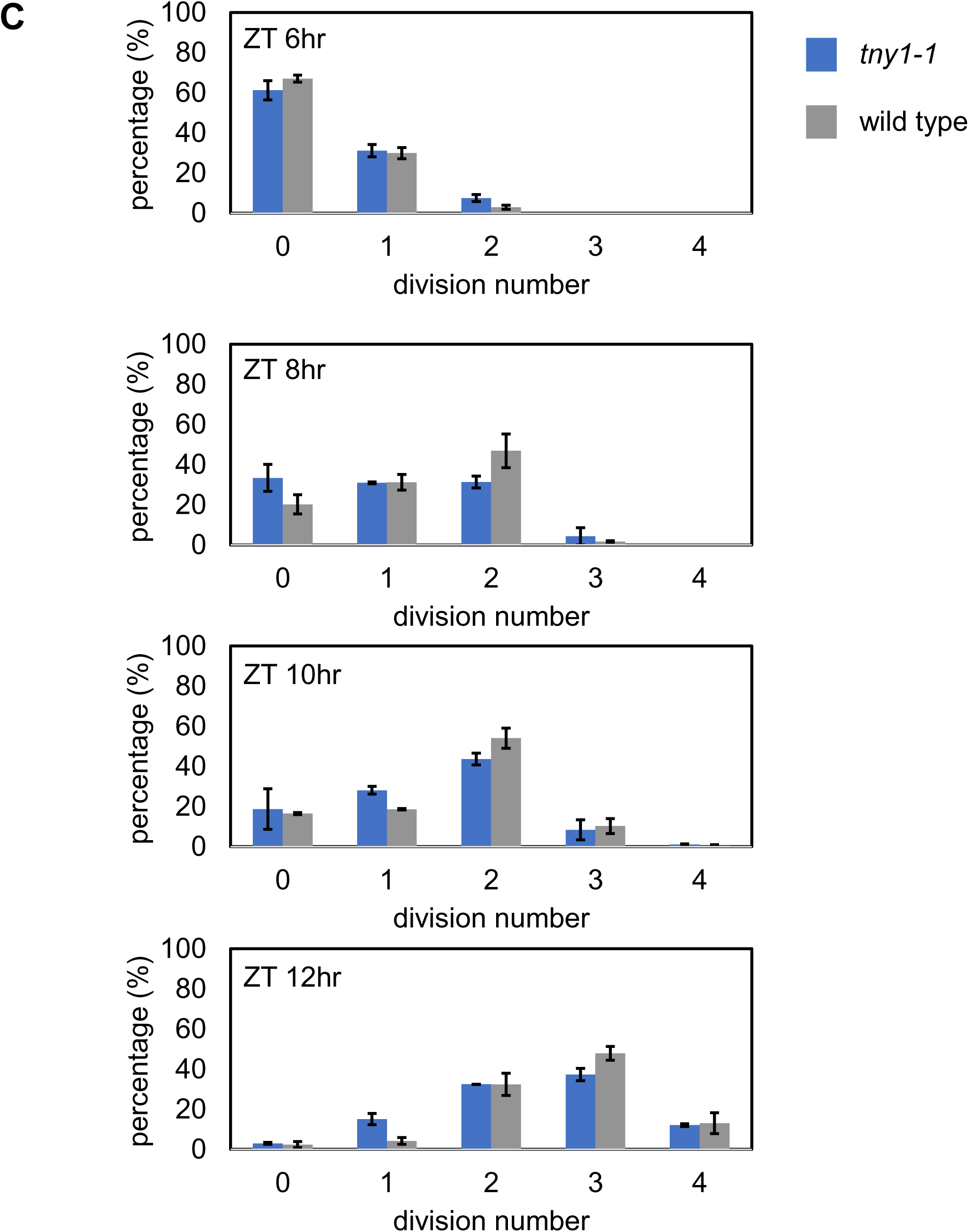
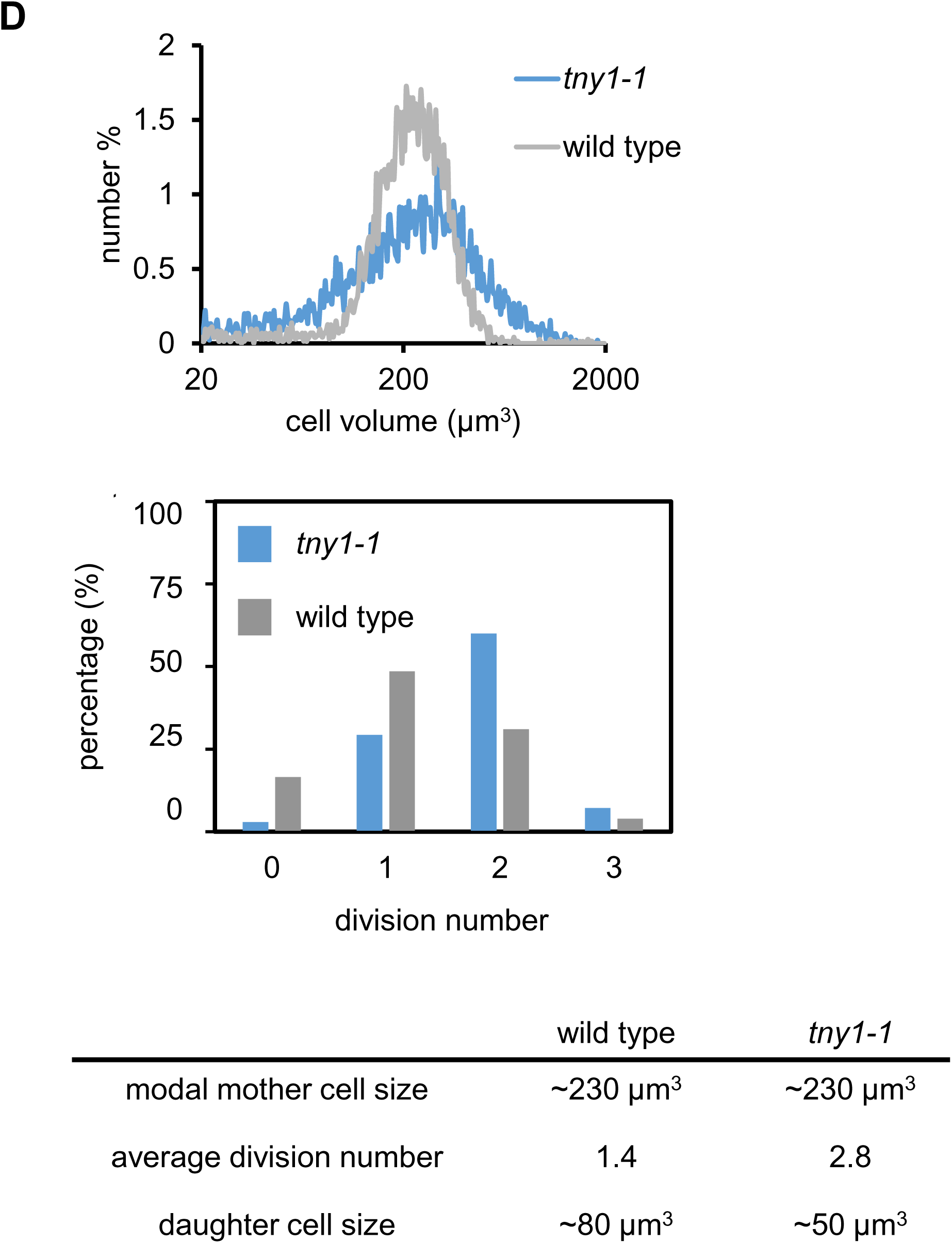
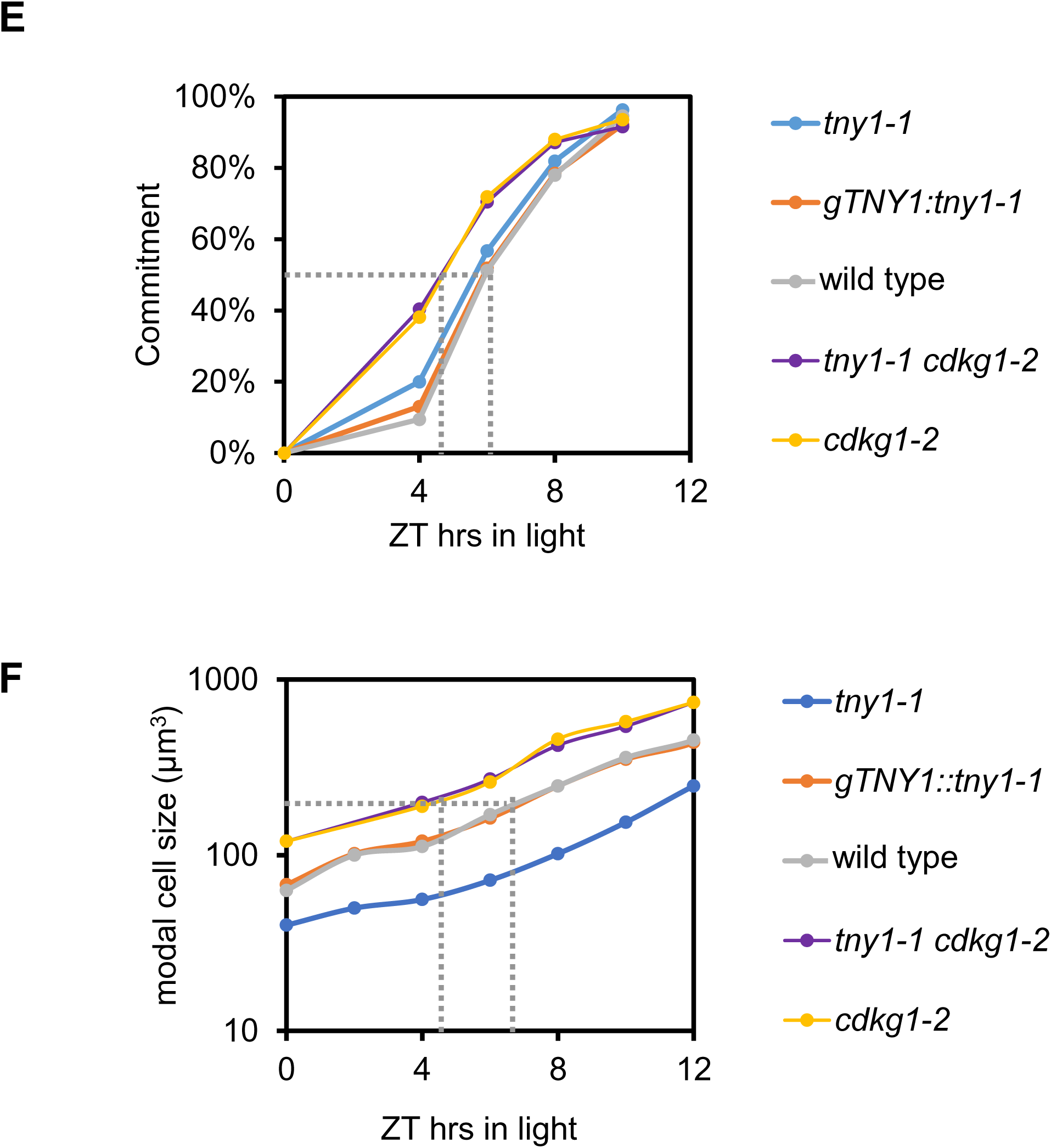
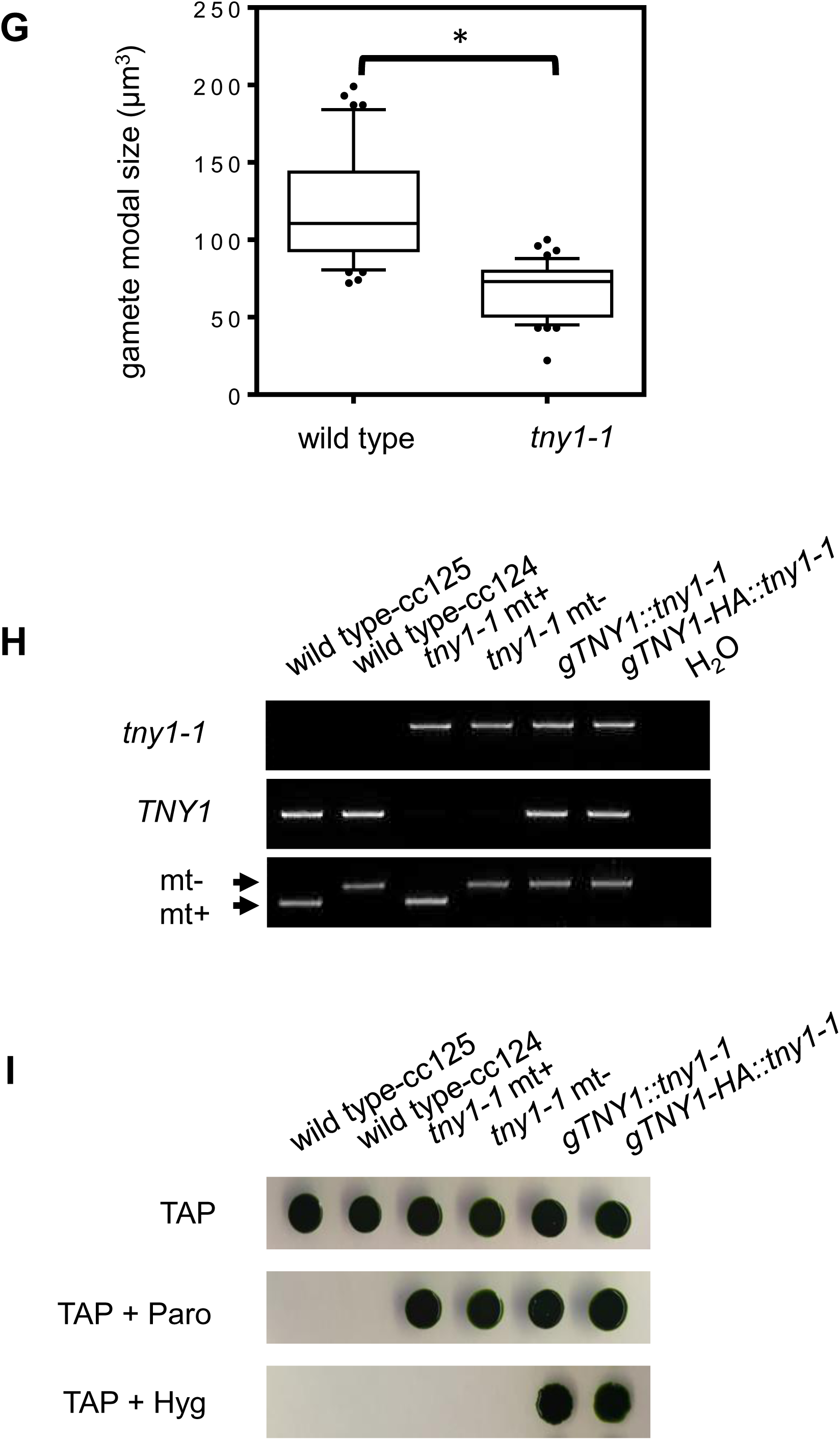
Characterization of *tny1-1* and rescued *tny1-1* strains. (A) Plot showing passage through Commitment (Commitment %, solid lines) and mitotic index (fraction dividing %, dashed lines) of synchronous *tny1-1*, wild type CC124, and a *tny1-1* rescued strain *gTNY1::tny1-1* collected at indicated time points during a synchronous diurnal cycle. Grey dotted line marks the time when 50% of the cells had passed Commitment (∼5 hrs ZT). (B) Plot of modal cell sizes for cultures in panel (A). Grey dotted line marks cell size at 50% Commitment. (C) Division number profiles of *tny1-1* and wild type CC124 from same experiment as in panel (B). At the indicated time cells were plated on minimal media, incubated in the dark, and scored for cell divisions (see Methods). (D) Division number profiles of size-matched G1 phase cultures of *tny1-1* and wild type cells (∼230 µm^3^) taken from different times in G1 to enable tny1-1 cultures to reach the same size as wild-type. Summary of results are in the table below. (E) Plot showing timing of Commitment for indicated genotypes, similar to panel (A). Grey dotted lines mark Commitment timing of *cdkg1-2* or *tny1-1 cdkg1-2* and wild type. (F) Plot of modal cell sizes for cultures in panel (E). Grey dotted lines mark cell sizes of strains in panel (E) showing that *cdkg1-2* and *tny1-1 cdkg1-2* have similar Commitment sizes as wild type. (G) Box and whisker plots of modal gamete sizes for individual wild type (n=44) or *tny1-1* (n=46) segregants from a back-cross between wild type and *tny1-1* (Methods). Boxes enclose the second quartile of data with horizontal lines showing median values, and whiskers enclose the 10^th^ - 90^th^ percentiles. Outliers are plotted as individual data points. Results of a Student’s t test are shown above (*, p<0.01). (H) Genotyping of the indicated strains for *tny1-1*, *TNY1*, and mating type loci (mating type minus, mt-; mating type plus, mt+). (I) Growth on selective media for *tny1-1* (paromomycin resistance marker; Paro) and *tny1-1* with rescuing constructs introduced with a hygromycin resistance marker, Hyg.

**Figure S2.**
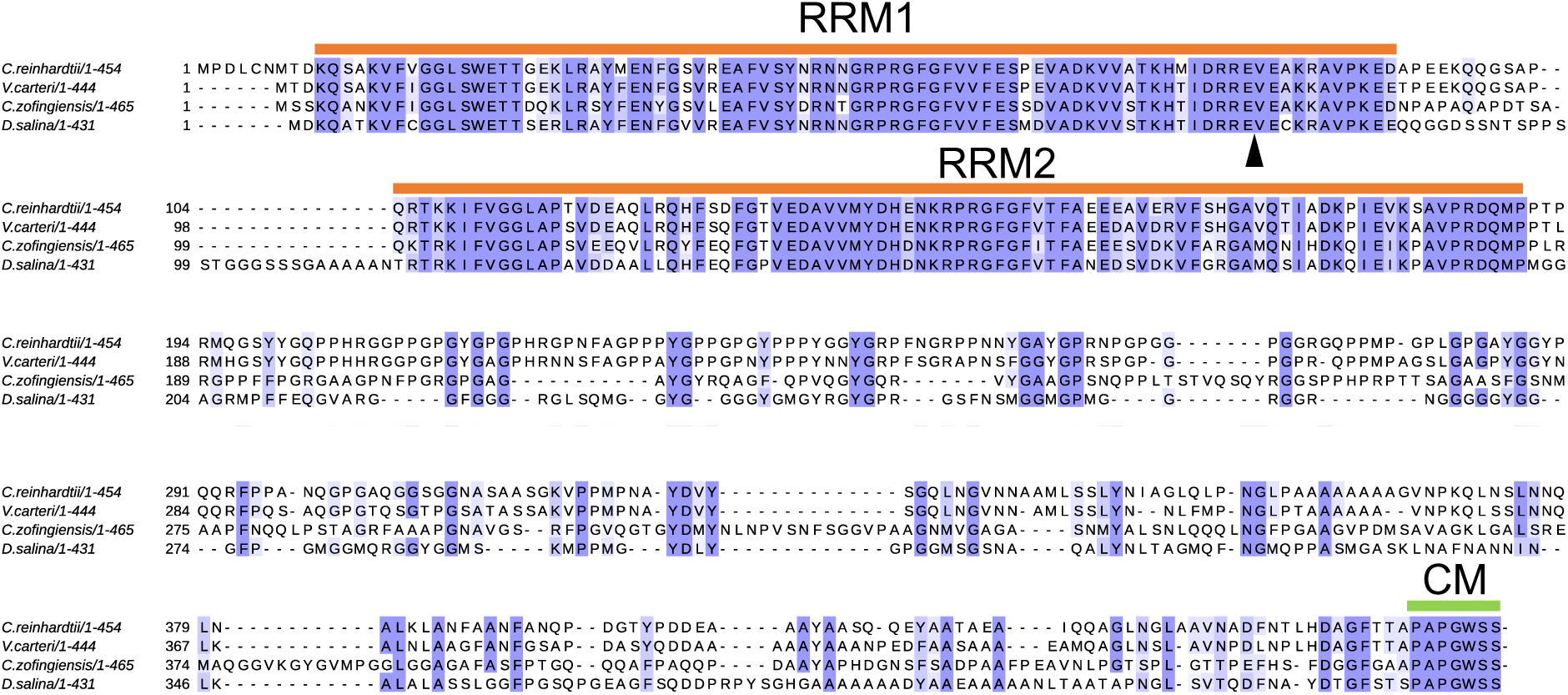
Multiple sequence alignment of green algal TNY1 orthologs. Peptide alignments for subset of proteins from Fig. 2: *Chlamydomonas reinhardtii* TNY1 (Cre07.g330300), *Volvox carteri* (Vocar.0031s0001), *Chromochloris zofingiensis* (Cz12g11070), and *Dunaliella salina* (Dusal.0065s00006). Gene IDs are from Phytozome (17). Alignment is shaded to show conserved residues. Positions of RNA recognition motifs 1 and 2 (RRM1, RRM2) and a conserved C-terminal motif (CM) are marked. The inverted black triangle shows the position the single intron found in subclade of algal TNY1 orthologs.

**Figure S3.**
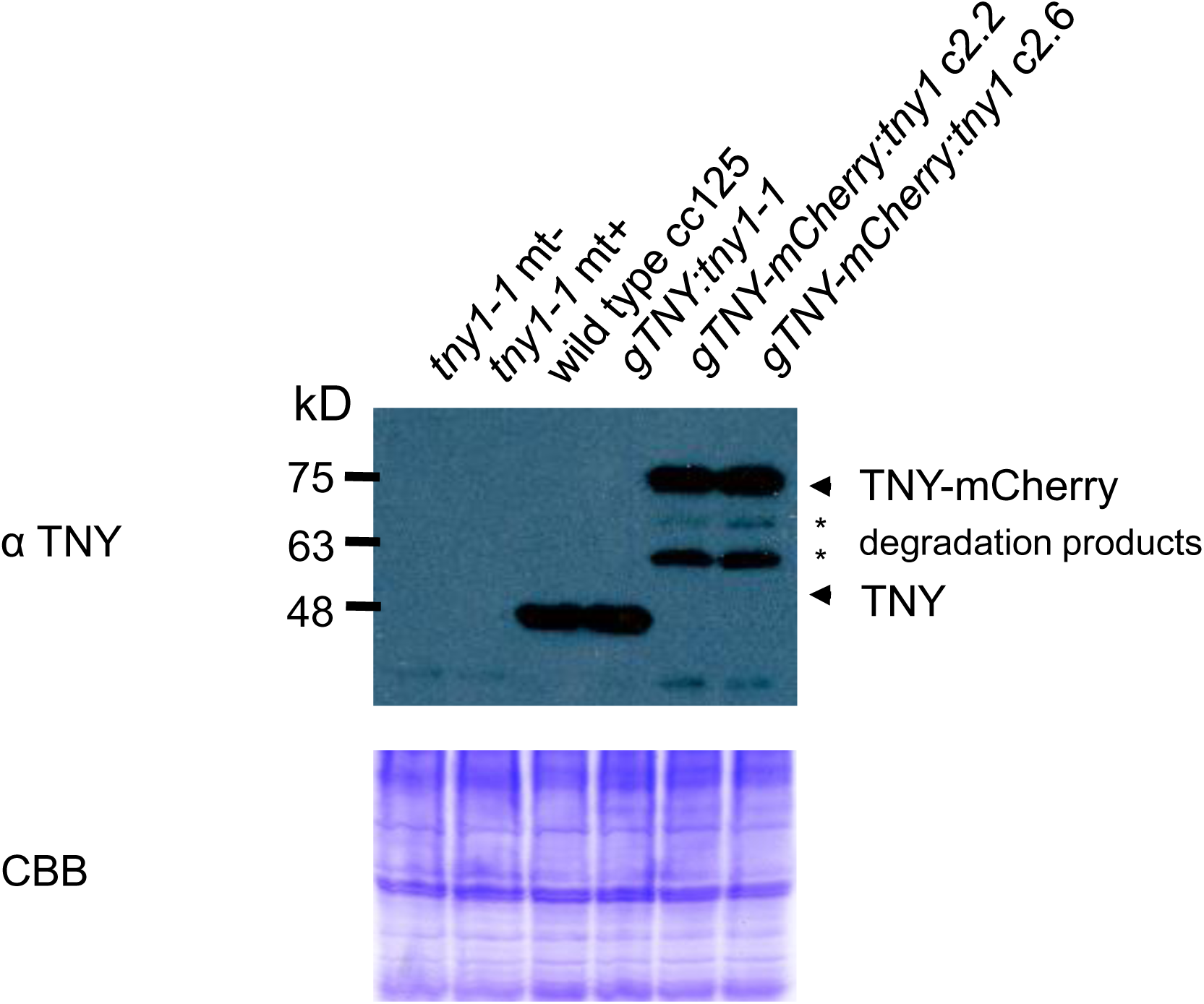
Detection of TNY1-mCherry expression in *gTNY-mCherry:tny1-1* strains. Whole cell lysates of daughter cells from indicated genotypes were loaded with equal biomass per lane, fractionated by SDS-PAGE, and immunoblotted using α-TNY1 (upper panel). Coomassie blue (CBB) staining is shown in the lower panel.

**Figure S4.**
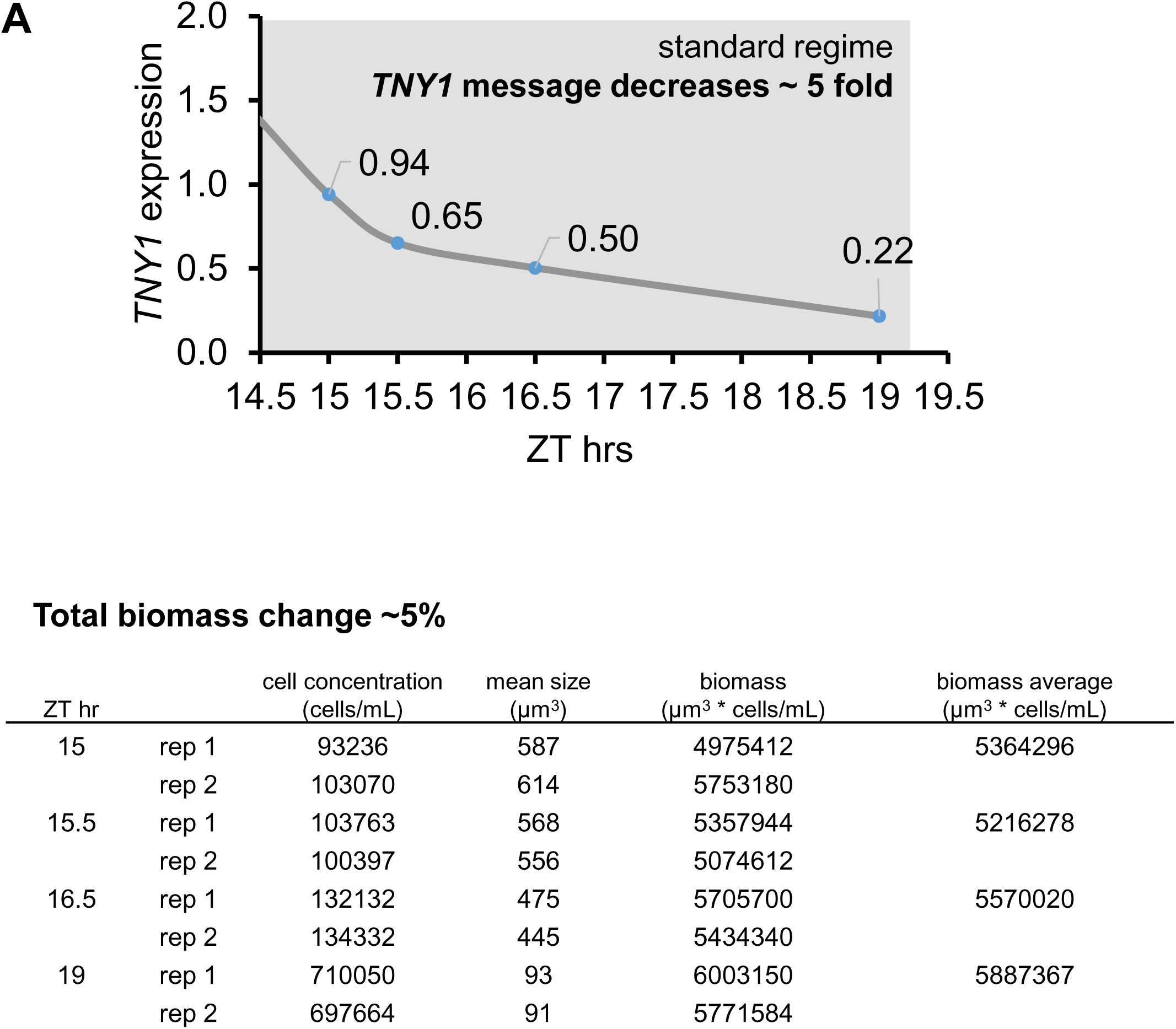
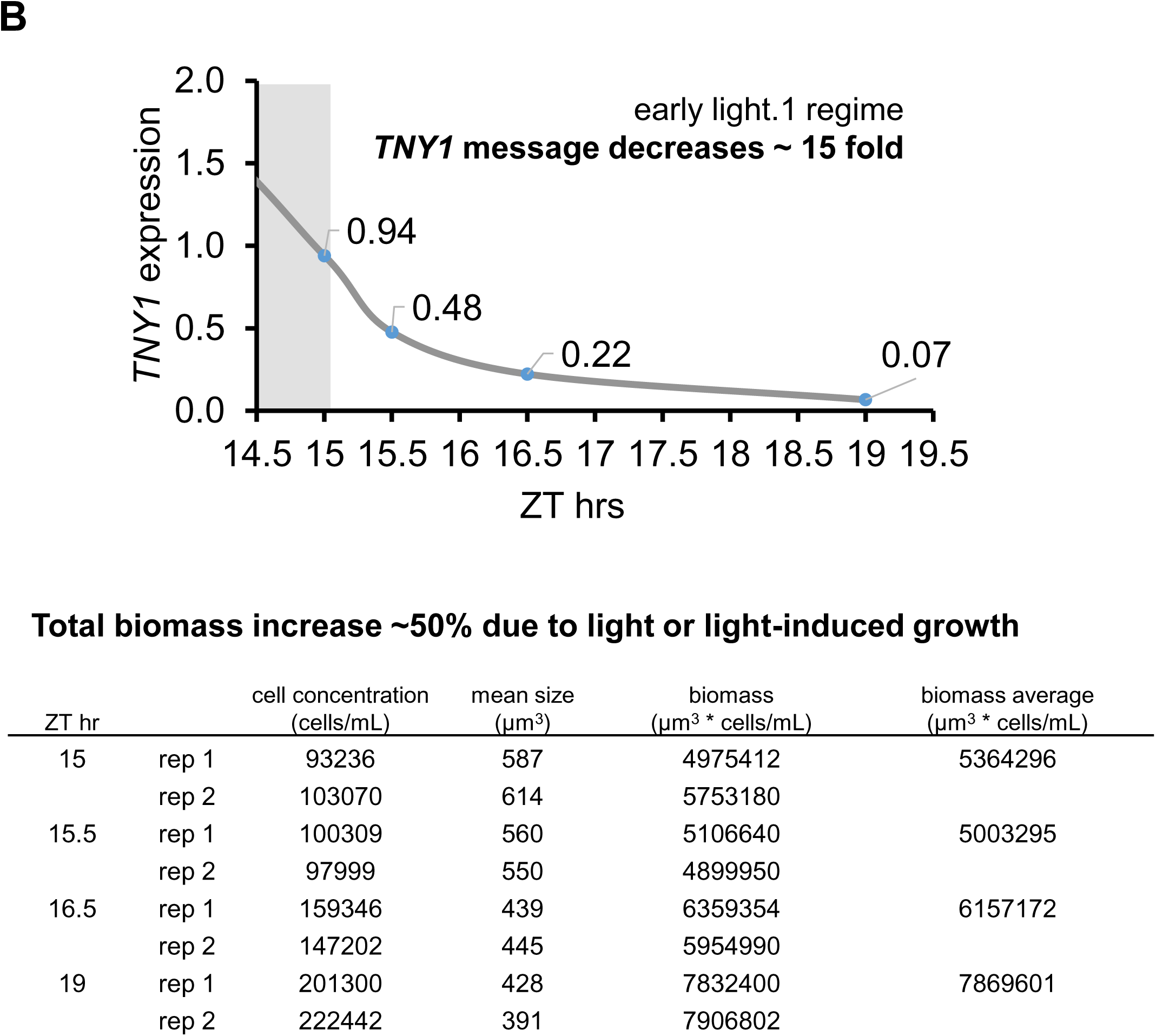
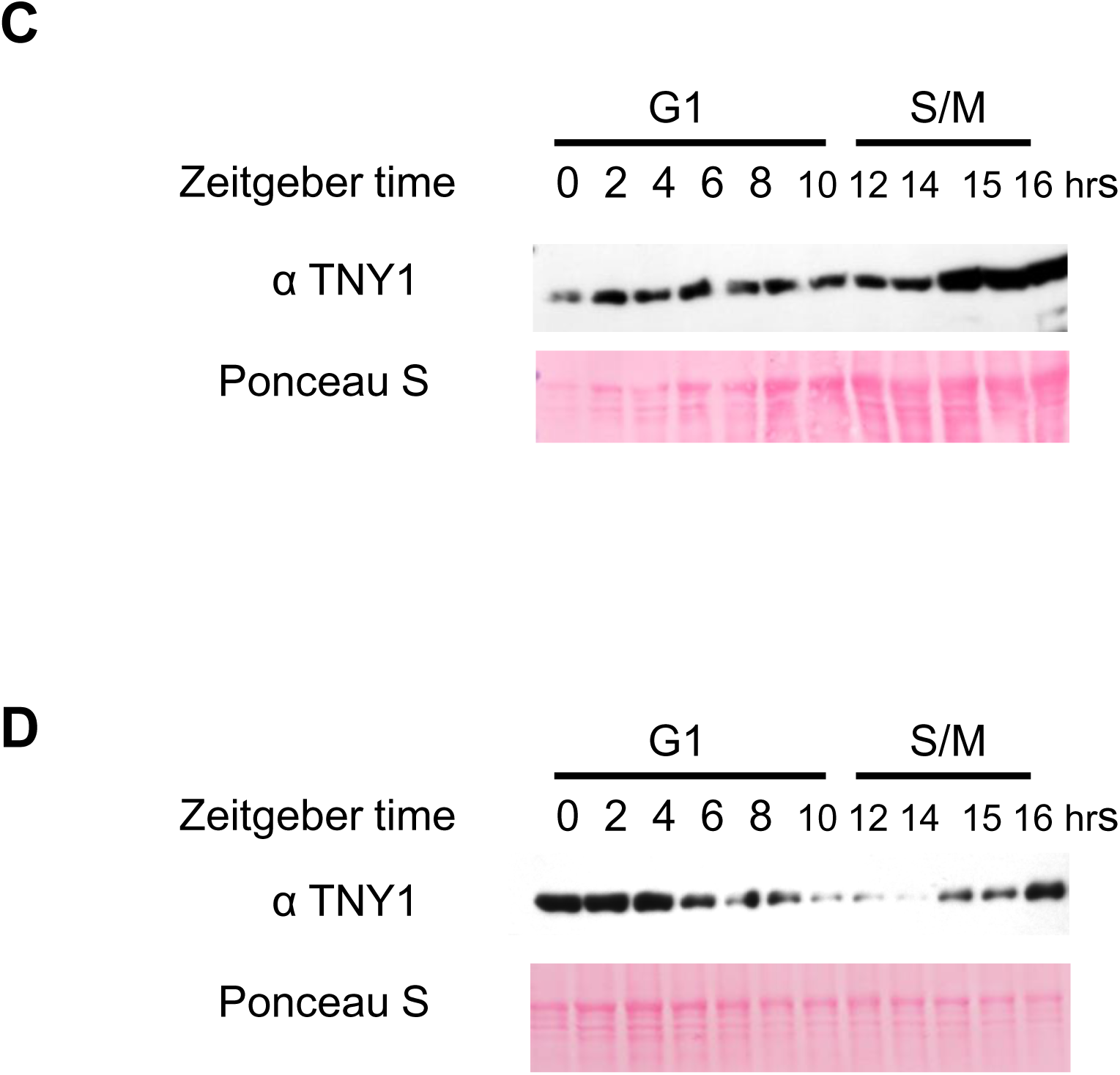
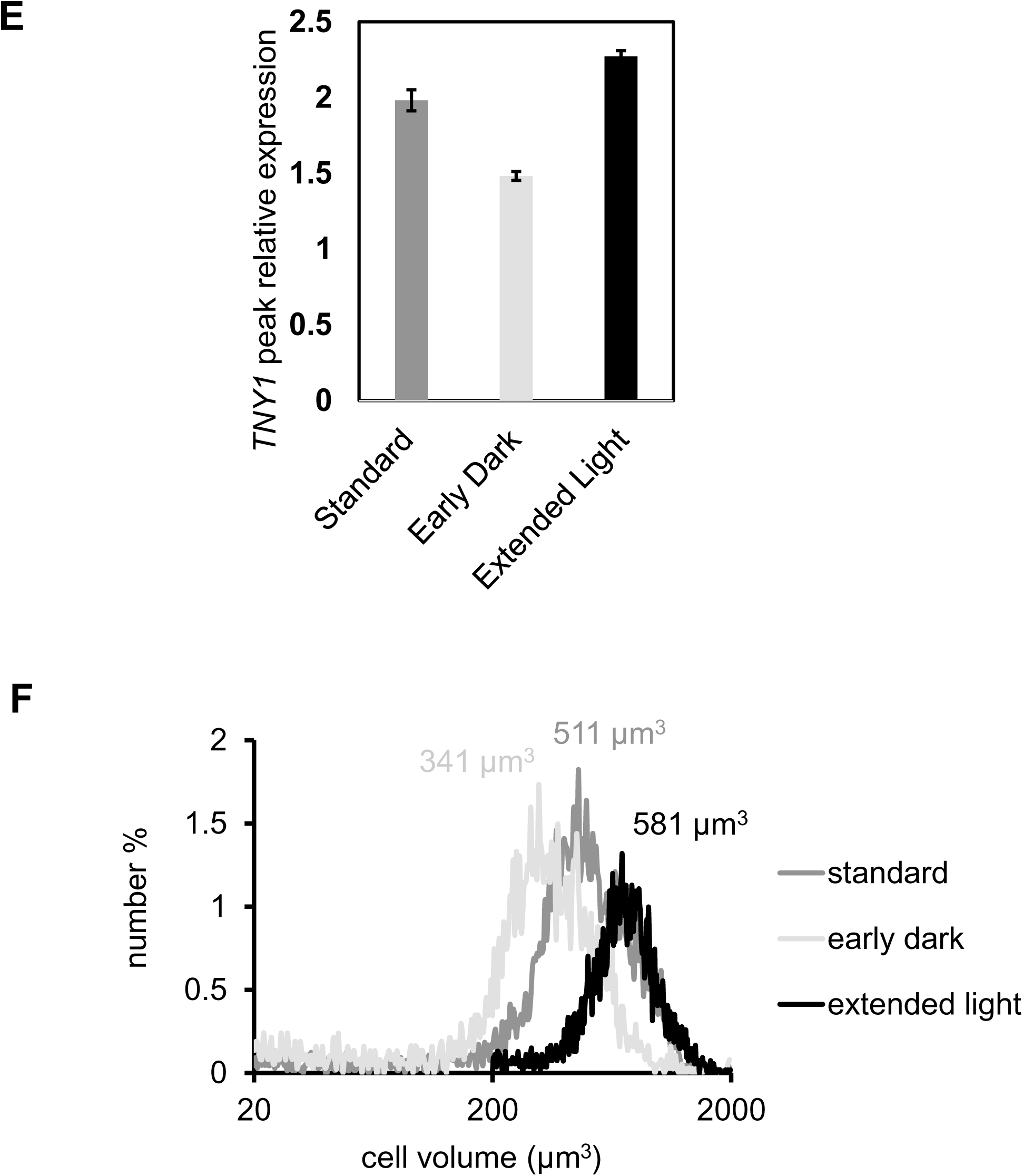
Cell cycle and diurnal control of *TNY1* mRNA and TNY1 protein accumulation. (A) and (B) qRT-PCR data time series for *TNY1* mRNA accumulation at ZT 15, 15.5, 16.5 and 19 in cultures similar to Figure 4A. (A), standard regime - 12hr:12hr light:dark. (B), early light1 regime with lights on at ZT 15. All data were normalized against control transcript *GBLP.* Error bars: high and low values of two biological replicates. The value of each biological replicate is calculated as the average of two technical replicates. The corresponding tables below each qRT-PCR data summarized the biomass change by comparing cell concentration (cells/mL) and mean cell size (µm^3^), and the calculated biomass (µm^3^ * cells/mL) of the two replicates at each ZT under the two regimes. (C) and (D) Biological replicates of immunoblots described in Figs. 4C and 4D. (E) *TNY1* mRNA accumulation peaks at ZT 15 under different diurnal regimes in Figure 4A. Error bars: high and low values of two biological replicates. The value of each biological replicate is calculated as the average of two technical replicates. (F) Size distributions of pre-division populations at ZT 15 under different diurnal regimes in Figure 4A. Numbers above the curves are the mean cell size of the corresponding populations.

**Figure S5.**
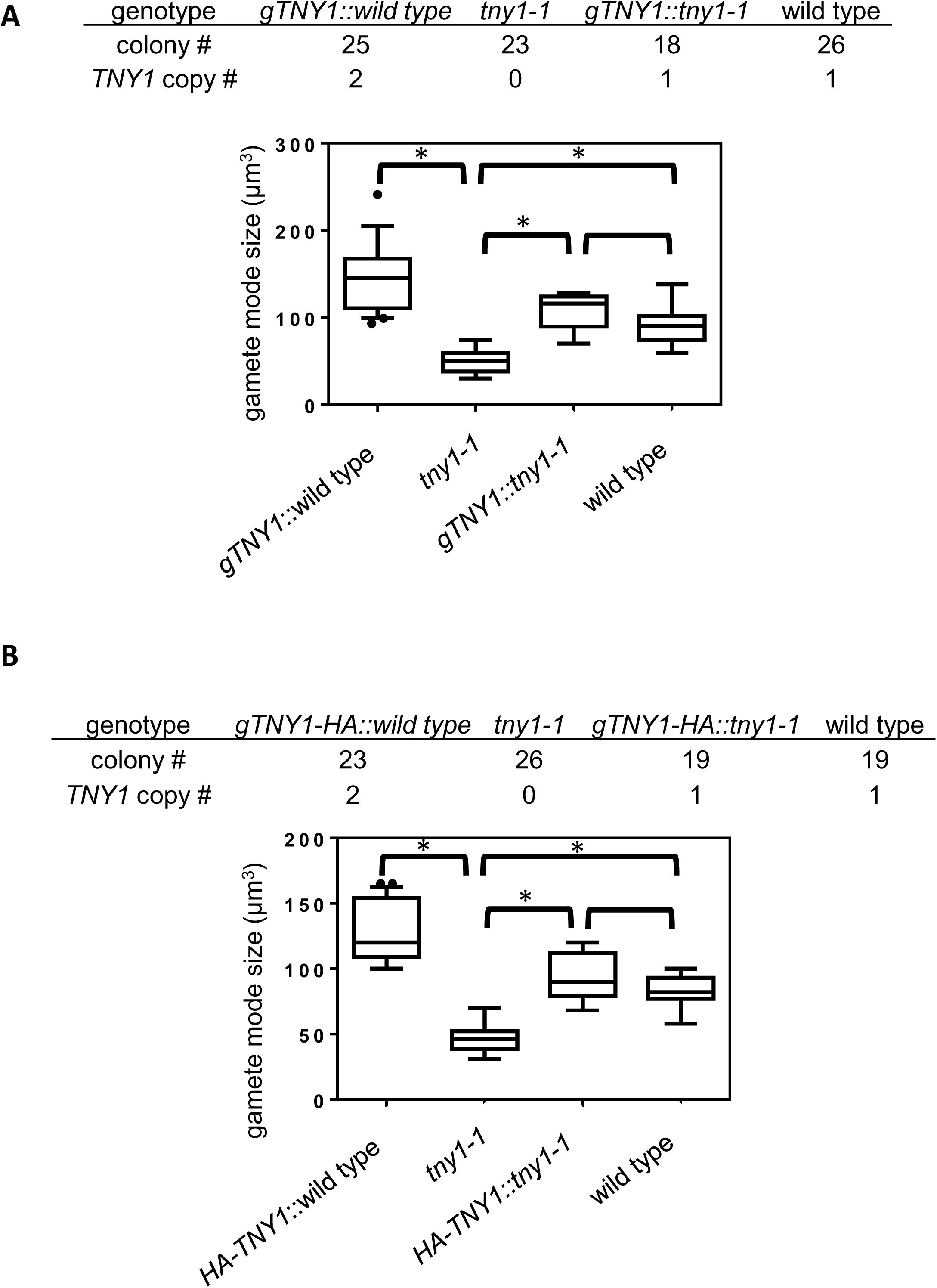
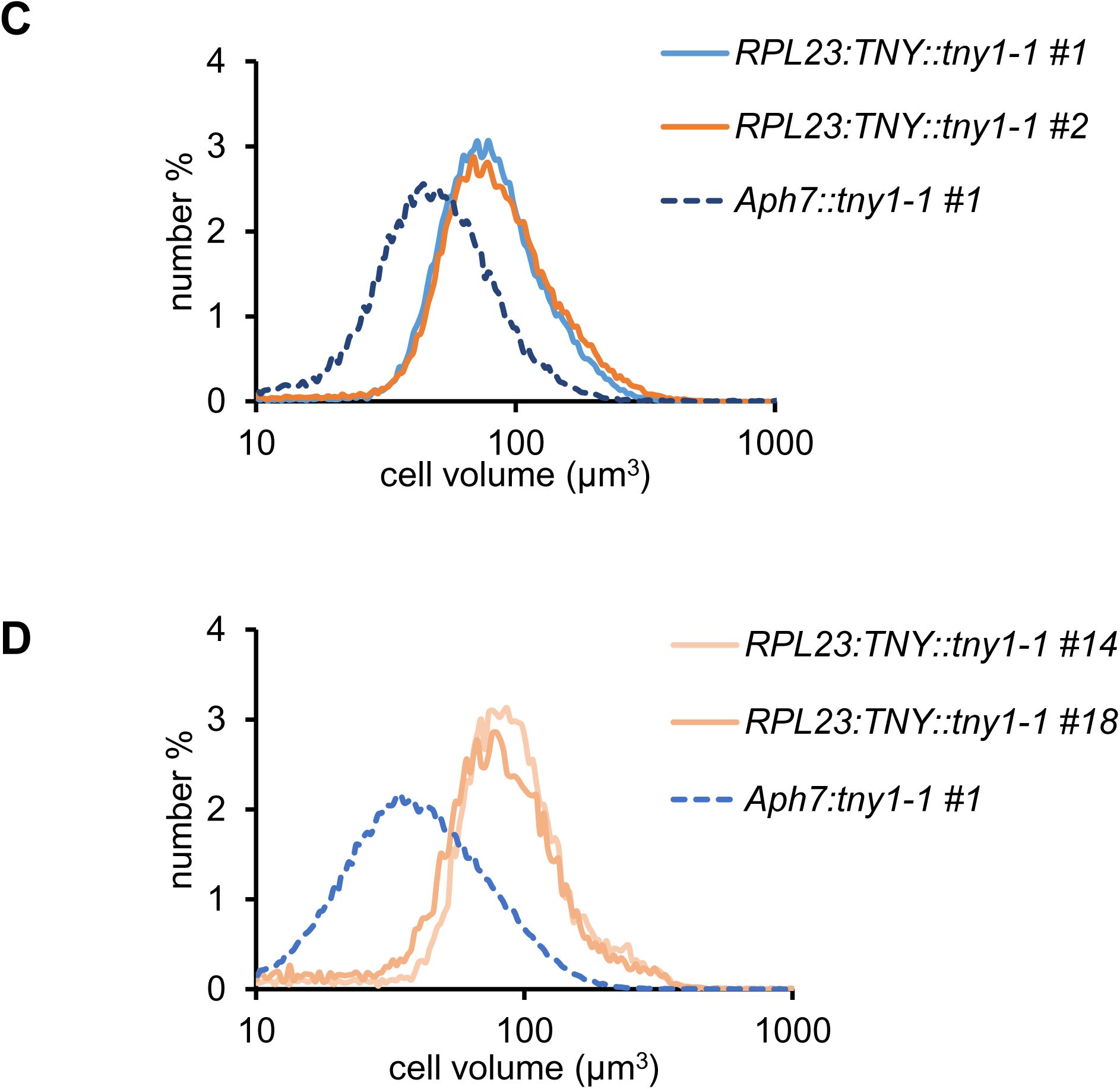
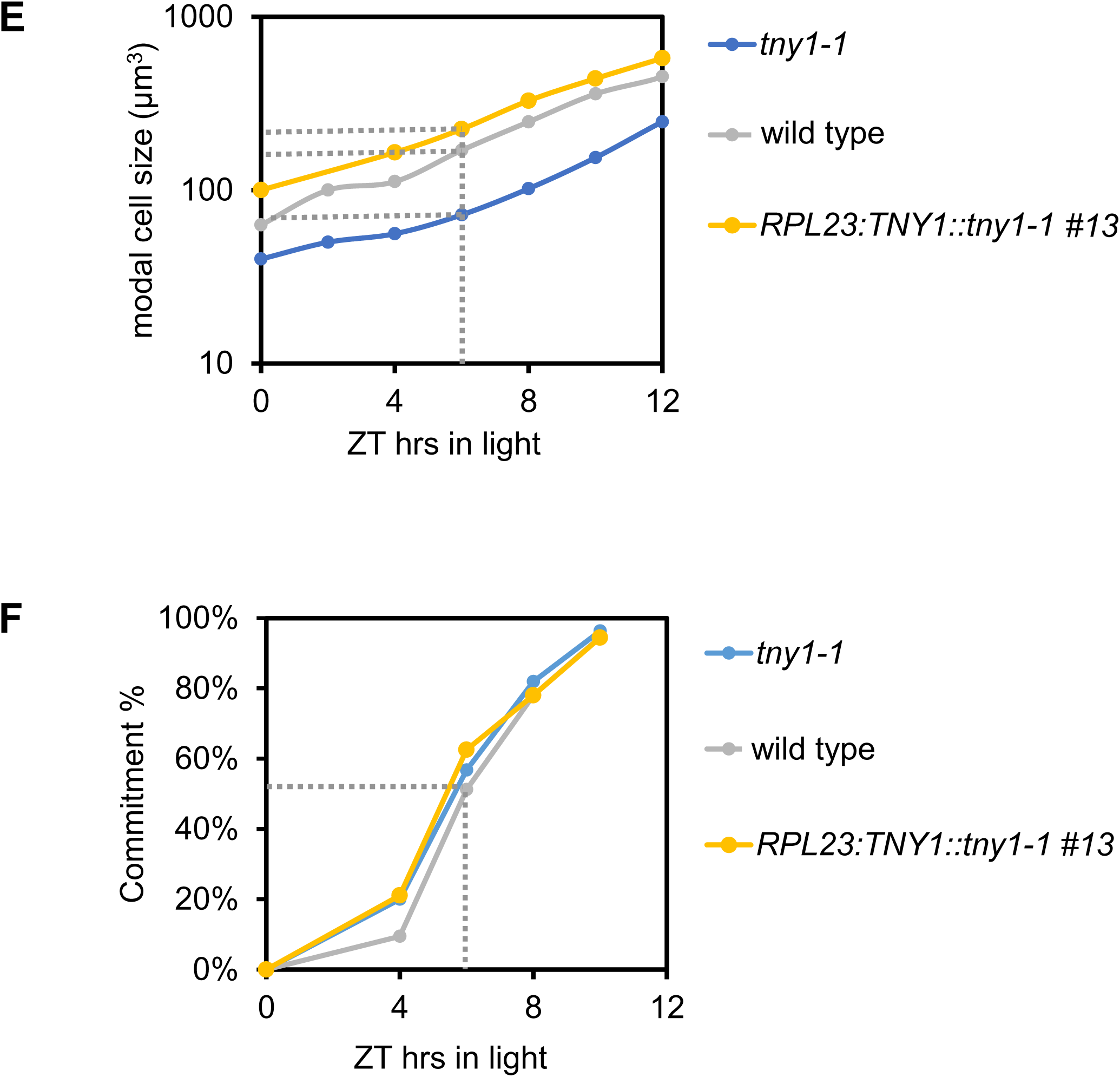
Dosage sensitivity of *TNY1* in haploid crosses. (A) Box and whiskers plots of modal gamete sizes for individual progenies from a back-cross between wild type CC124 and rescued strain *gTNY tny1-1.* with genotypes *gTNY1 TNY1* (n = 25), *tny1-1* (n =23), *gTNY1 tny1- 1* (n = 18), and wild type (n = 26). Boxes enclose the second quartile of data with horizontal lines showing median values, and whiskers enclose the 10^th^ -90^th^ percentiles. Outliers are plotted as individual data points. Results of a Student’s t test are shown above (*, p<0.01). (B) Similar data to (A) for a back-cross between wild type CC124 and *HA-TNY tny1-1*. (C) Size distributions of daughters from representative independent *RPL23:TNY tny1-1* strains which show rescued cell size (modal size ∼80 µm^3^) and a control transformant with madeusing an empty vector *Aph7 tny1-1* (modal size ∼50 µm^3^). (D) Size distributions of daughters from independent *RPL23:TNY tny1-1* strains which show a large-size phenotype (modal size >100 µm^3^) and a control strain *Aph7 tny1-1* (modal size ∼50 µm^3^).

**Figure S6.**
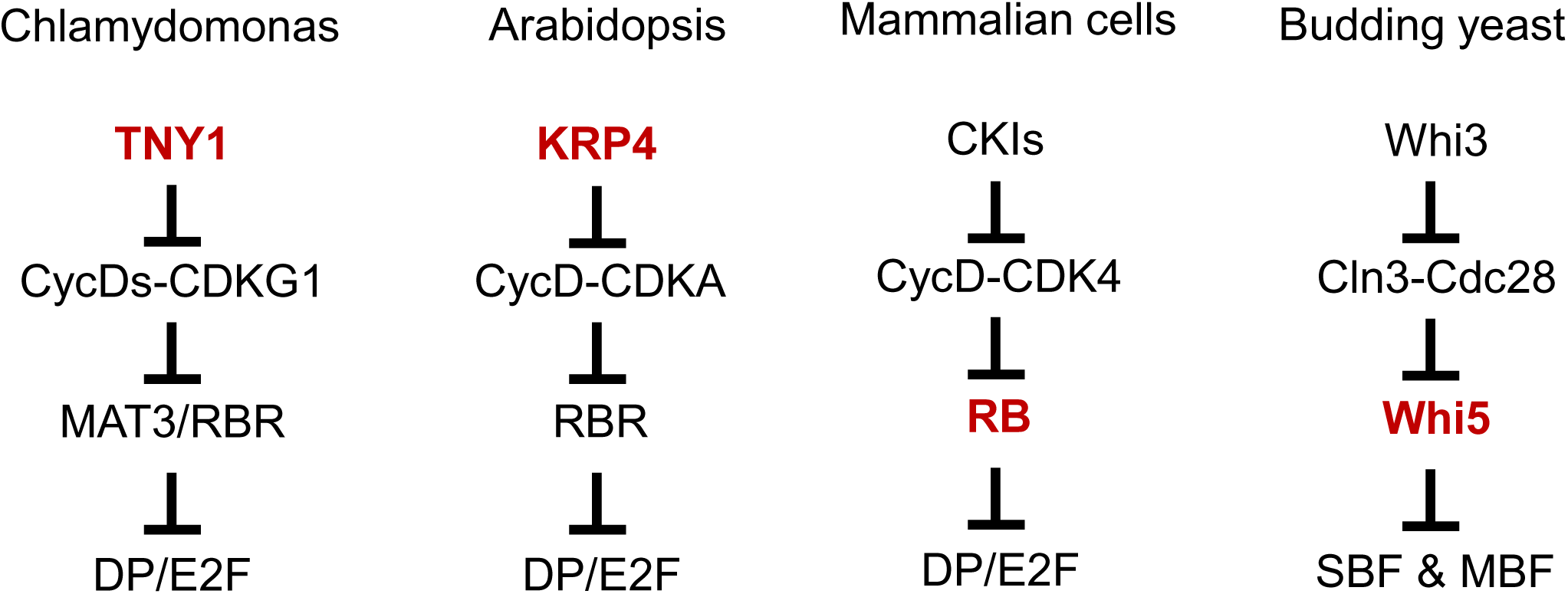
Systems level comparison of cell size control across taxa. Cell cycle inhibitors subscaling with cell size in G1 phase are in bold red.

**Table S.**
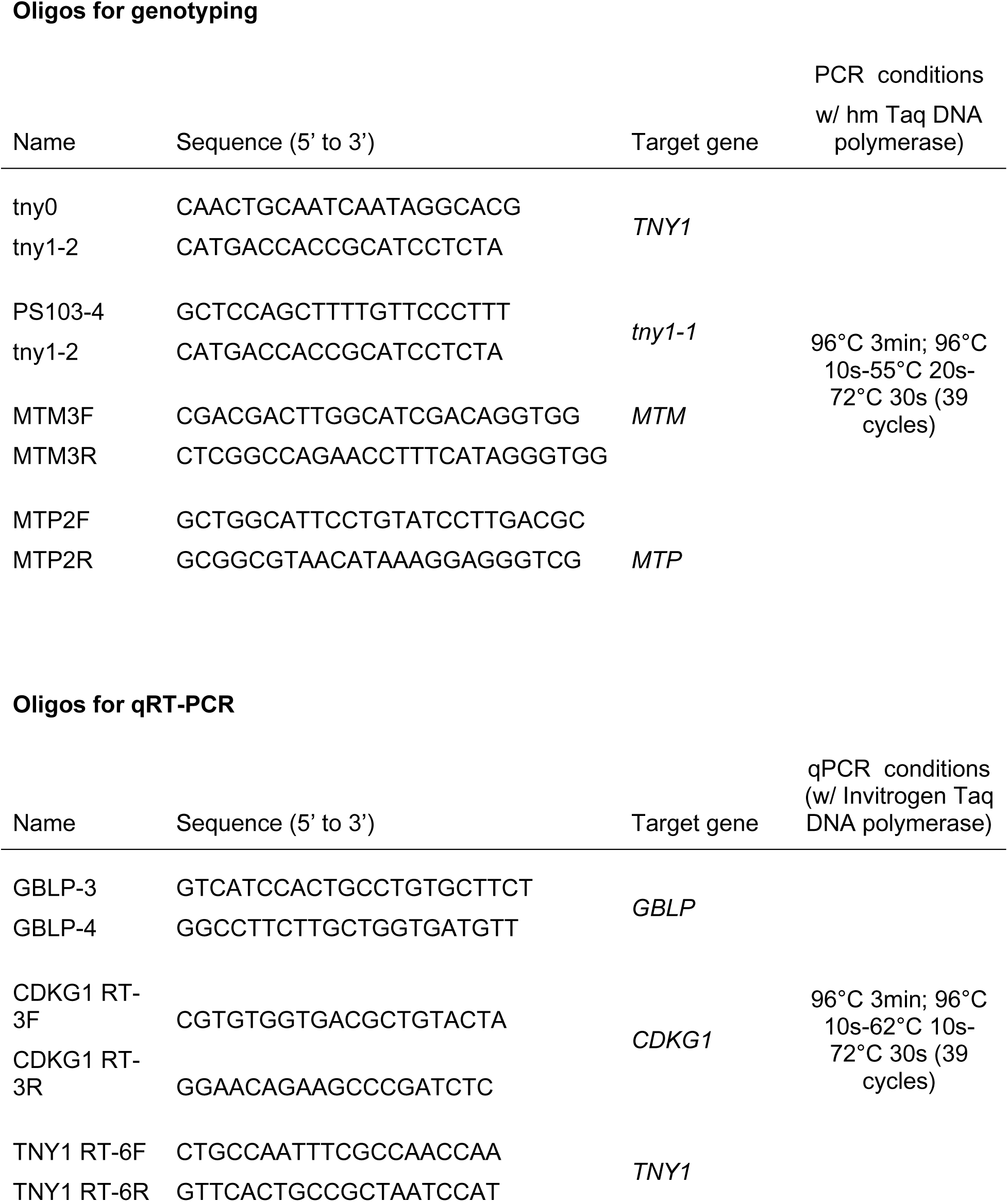
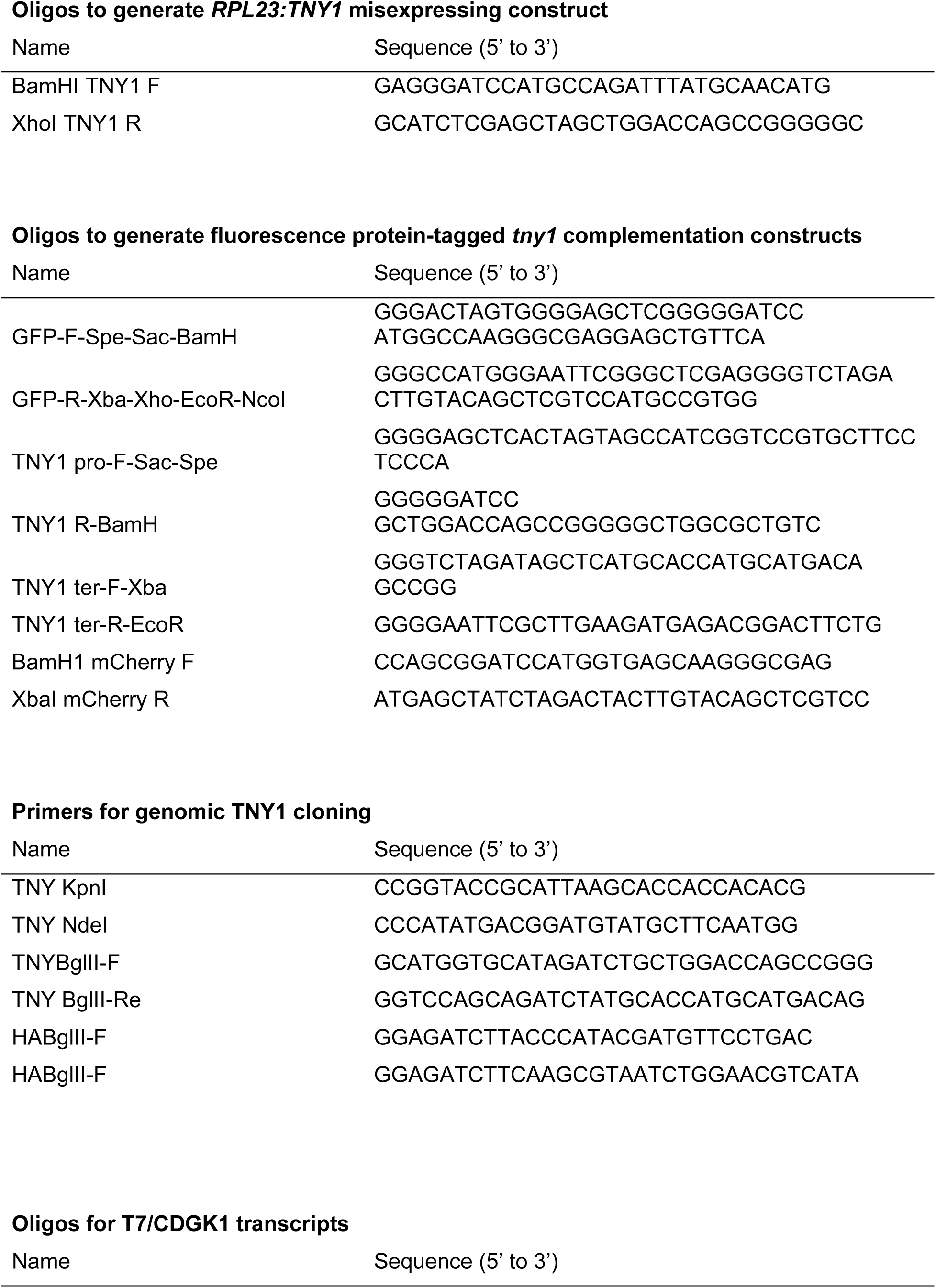
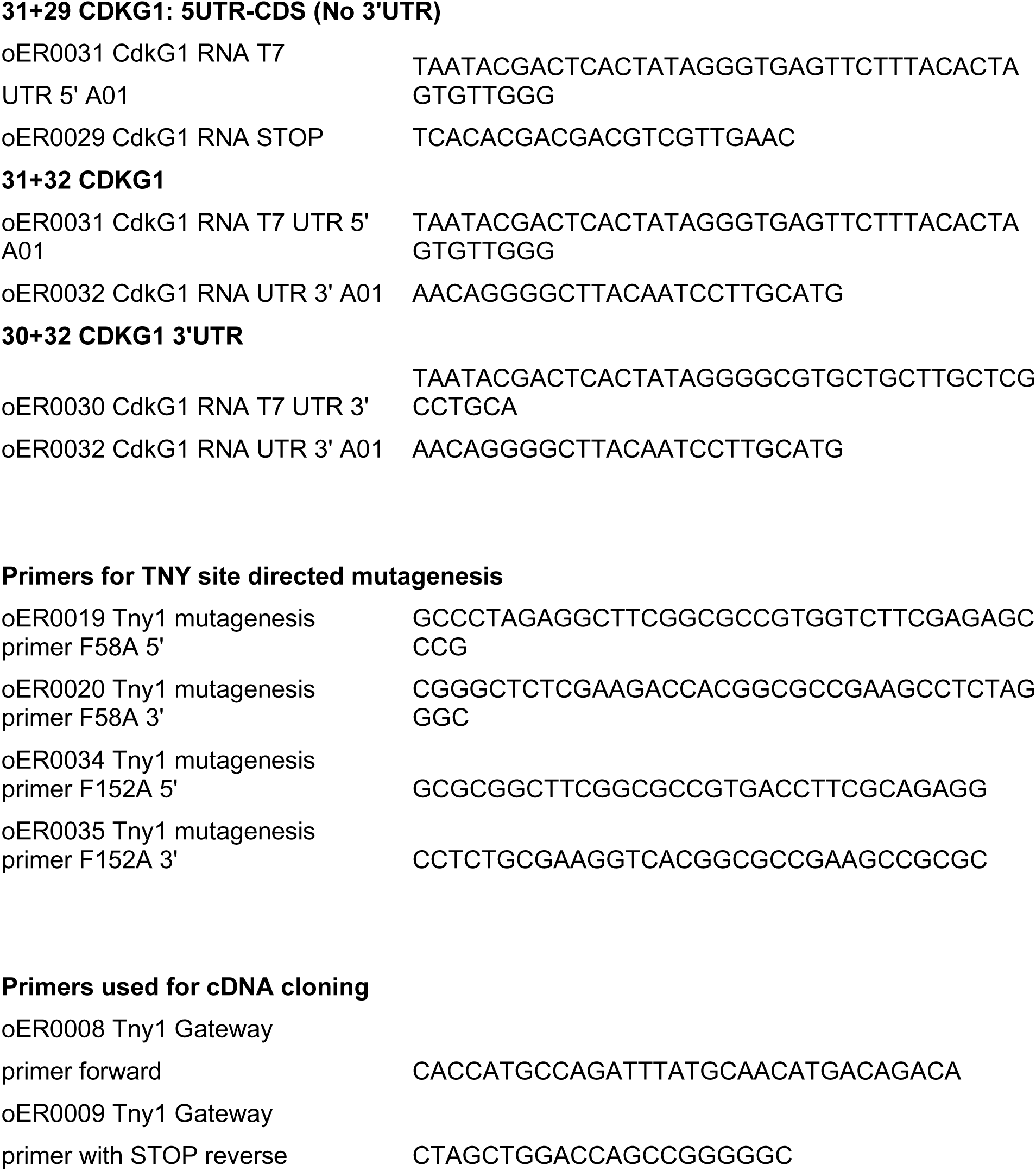
Oligos used in the study.

